# Virome DNA stable isotope probing reveals diverse active soil virus communities across ecosystem contexts

**DOI:** 10.64898/2026.03.02.709135

**Authors:** Ernest D Osburn, Minjae Kim

## Abstract

Viruses are increasingly recognized as important players in soil ecosystems, but the active lytic virus populations that influence microbe-mediated terrestrial ecosystem processes remain mostly uncharacterized. Here, we trace ^13^C-labelled glucose from host microorganisms into virus genomic DNA to identify virus populations actively involved in soil carbon (C) cycling, *i.e.*, viruses that lysed ^13^C-incorporating microbes. We present experimental evidence of isotope labelling (*i.e.*, lytic activity) of more than 5,000 virus populations. The active viruses lysed hosts from 197 microbial families across 28 prokaryote phyla. Viral lysis was greater in C-limited agricultural soils compared with C-rich forest soils, highlighting C availability/inputs as key factors that mediate virus life cycles. Active viruses disproportionately lysed microorganisms in the Bacillota, Bacteroidota, and Pseudomonadota phyla, likely reflecting glucose-induced growth responses of microbial copiotrophs within those groups. Supporting this, we observed that the degree of virus genome isotope labelling was positively correlated with the growth potential of the microbial hosts. Furthermore, the active viruses exhibited unique genomic characteristics compared to the inactive viruses, including greater prevalence of lysogeny-associated genes and distinct profiles of putative auxiliary metabolism genes in the active virus genomes. Overall, our results demonstrate a link between microbial growth traits and virus activity and suggest that substrate-induced viral lysis significantly influences microbial population turnover in soil. Our results also show that virus activity in response to C inputs is highly variable among soil contexts, with implications for the varying ecosystem-scale influences of viruses among terrestrial environments.

## Introduction

Viruses are ubiquitous components of natural environments, influencing population-, community- and ecosystem-scale ecological processes in a variety of habitat types. Of particular note are viruses that infect prokaryotes, which are the most abundant biological entities on earth (1, 2) and which influence the abundance, diversity, and metabolic functions of their microbial hosts (3–5). However, the ecological roles and importance of viruses in soil remain very poorly understood (6–9), in contrast to better-characterized environments such as marine waters (10, 11). This is despite the vast abundance of viruses in soil, where viruses typically outnumber their cellular hosts more than 10-fold (1, 2) and where 30-100% of microorganisms are infected with viruses at any given time (12, 13). Ultimately, our poor understanding of soil virus ecology limits our understanding of microbe-mediated terrestrial ecosystem functioning, which highlights the importance of expanding our knowledge of the complex virus-microbe interactions in soil.

Facilitated by advancements in virus isolation methods and sequencing/bioinformatics technologies (14–17), soil virus ecology research has begun to accelerate in recent years (6, 8, 18). A central theme that has emerged from this research is that soil virus communities strongly covary with soil physicochemical properties; for example, soil C and nitrogen (N) content have been shown to be important explanatory factors associated with biogeographic patterns in soil virus communities (19, 20). Supporting this finding, other studies have demonstrated responses of virus communities to soil N additions (21, 22) and to the C:N ratios of soil amendments (23). Furthermore, genomic analyses have shown that soil viruses contain auxiliary metabolism genes (AMGs) that potentially influence the C and N cycle functions of their microbial hosts (17, 24). Taken together, these studies suggest that soil virus communities both respond to and potentially influence C and N cycling in soil. To investigate the mechanisms underlying these patterns, a parallel line of inquiry has emerged using experimental manipulations of soil viruses to quantify the effects of viruses on terrestrial ecosystem processes. These studies have demonstrated influences of viruses on soil N turnover (25), litter decomposition rates (26), and soil C cycling (27–30). However, the soil ecosystem responses to virus manipulation that have been detected vary in direction and magnitude among studies, leading to uncertainty regarding the functions and importance of viruses in soil. For example, viruses have been shown to increase, decrease, or not change soil C mineralization rates, depending on the study in question (26–30). These differences likely originate from differences among soil and experimental contexts in which viral influences were quantified and highlight our lack of fundamental understanding of how virus-microbe interactions vary across soil contexts.

A key unresolved question regarding virus-microbe interactions in soil is the degree to which viruses influence microbial community composition and turnover rates through lytic replication (31). One mechanism by which these influences could manifest is the “cull the winner” model of virus replication, where viruses selectively lyse successful, rapidly growing microbial populations in a density-dependent manner (32). These lysis events could influence both microbial community structure and soil biogeochemical cycles by accelerating microbial population/biomass turnover (27). However, other models of virus-microbe interactions have also been proposed, *e.g*., the “piggyback the winner” model where viruses favor lysogeny when their microbial hosts experience population booms (33, 34) and the “king of the mountain” model, which predicts that abundant microbial populations have an advantage in evolving resistance to viruses (35). Under these two alternative models, viral influences on soil ecological processes would be largely indirect and more difficult to isolate and quantify. It is important to evaluate support for these alternative models of virus-microbe interactions under varying soil environmental conditions given that the different mechanisms likely exist simultaneously in soil (34) and that their relative importance has implications for microbe-mediated terrestrial ecosystem processes (35).

One possible approach for identifying and quantifying virus lytic replication in soil is stable isotope probing (SIP), a technique that has been used to identify active cellular microbial populations by detecting the incorporation of heavy isotopes into microbial genetic material (36). This may also work to identify active virus populations in the lytic cycle since the elements that comprise virions are derived from the host organism. Indeed, some studies have successfully demonstrated this by recovering virus genomes from isotopically labelled total metagenomic DNA extracted from soil (4, 37–39). In so doing, these studies have identified specific virus populations with direct involvement in soil biogeochemistry, *e.g.*, the C and N cycles. However, these studies are also limited in their ability to identify active virus populations given the difficulty of assembling virus genomes from soil metagenomes due to the dominant presence of microbial DNA in total metagenomes (32, 40). In response to this limitation, one recent study attempted a direct approach to identifying active viruses by tracing ^18^O from labelled water into isolated virome DNA from soil (41). That study was able to identify several dozen active virus populations, mostly linked to microbial hosts in the Actinomycetota and Pseudomonadota phyla. Developing and extending approaches such as these will be critical for expanding our knowledge of virus activity in soil.

In this study, we employed a novel DNA SIP method in which we traced ^13^C-labelled glucose into isolated virome DNA to characterize virus populations that lyse their microbial hosts following labile C inputs to soil. We conducted the isotope probing in soils from contrasting ecosystems (*i.e.*, agriculture, forest) to explore how environmental context influences substrate-dependent virus activity. By implementing this method, we sought to assess relationships between host microorganism characteristics and the degree of lytic viral activity in the context of labile C additions. Specifically, we hypothesized that virus activity in response to glucose addition would be greatest in strictly lytic populations that infect highly copiotrophic microbial lineages, reminiscent of the “cull-the-winner” model (32). Related to this, we also predicted that variation in lytic virus activity between the soil types would be attributed to differences in overall growth potential of the resident microbiomes in the contrasting soil environments. Alternatively, if the “piggyback the winner” or “king of the mountain” models of virus-microbe interactions predominate, we might observe little evidence of increased viral lysis in response to labile C additions to soil. Our study provides new insights into the active soil virus populations that contribute to microbial turnover in soil and illustrates the benefit of combining SIP and viromics for interrogating virus-microbe interactions related to soil C cycling.

## Materials and Methods

### Site descriptions, soil sampling, and soil analyses

To characterize glucose-responsive active soil virus communities, we sampled soils from two sites across a land use intensity gradient: a highly disturbed agricultural field and an undisturbed forest ecosystem. The large variation in soil properties between the sites (see below) allowed us to explore how soil environmental context influences substrate-induced virus activity. While the two ecosystem types are represented by only one site each, the soils therein are representative of the common differences in physicochemical properties of the land uses, *e.g.*, C-vs. N-limitation in the agricultural vs. forest soils, respectively. The agricultural soils were from a long-term experiment at the University of Kentucky North Farm site (38° 07′ N, 84° 29′ W) with continuous maize production and annual soil tillage and N fertilizer application (336 kg/ha) since 1970 (42). The forest soils were collected from an experimental watershed at the University of Kentucky Robinson Forest (37° 32’ N 83° 20.8’ W) that has remained undisturbed for over 100 years (43). We collected five 10 cm depth mineral soil cores from four replicate sampling locations in each site using a 5 cm diameter corer and composited the five subsamples from each location, for a total of four composite soil samples from each site. At the agricultural site, the samples came from four replicate experimental plots 5.8 m × 6.1 m in size. In the forest site, the samples came from four sampling locations equivalent in size to the agricultural experimental plots, each located 50 m apart along a 200 m transect. In the forest site, we removed O-horizon material prior to soil collection. The four samples from each site were maintained as separate biological replicates throughout the experiment. Samples were returned to the lab on ice the same day and then sieved (2 mm), homogenized, and subsamples stored at -80°C or air dried. The remainder of the sample was stored at 4°C prior to the incubation experiments, which occurred within two weeks of sample collection.

Prior to the start of the experiment, we measured several physicochemical properties of the soils: pH, total C and N, microbial biomass C, extractable N, microbial basal respiration, gravimetric water content, and water holding capacity. Details of these measurements are provided in the Supplementary Methods. Soil physicochemical data are provided on Table 1. We also characterized prokaryote communities in the soils using amplicon sequencing of the V4-V5 region of the 16S rRNA gene. Details on DNA extraction, amplicon library prep, sequencing, and bioinformatics are provided in the Supplementary Methods.

**Table 1:** Physicochemical properties of the agricultural and forest soils. Values are means followed by one standard deviation in parentheses. Asterisks indicate significantly higher values from one-way ANOVA at the following significance levels: \*\**p* < 0.01, \*\*\**p* < 0.001.

| Variable | Ag. soils | Forest soils |
| --- | --- | --- |
| Soil pH | 6.63 (0.16)** | 5.29 (0.50) |
| Water holding capacity (g H<sub>2</sub>O g soil<sup>-1</sup>) | 0.51 (0.005) | 0.85 (0.109)*** |
| Total C (mg C g soil<sup>-1</sup>) | 13.7 (0.69) | 39.2 (13.4)** |
| Total N (mg N g soil<sup>-1</sup>) | 2.51 (0.59) | 2.91 (1.07) |
| Soil C:N | 5.69 (1.41) | 13.5 (1.03)*** |
| Extractable N (µg N g soil<sup>-1</sup>) | 1026 (376)** | 17.7 (2.91) |
| Microbial Biomass C (µg C g soil<sup>-1</sup>) | 91.0 (28.6) | 793 (154)*** |
| Basal respiration (µg CO<sub>2</sub>-C g soil<sup>-1</sup> h<sup>-1</sup>) | 0.39 (0.11) | 1.08 (0.25)** |
| Glucose response (fold-increase in respiration) | 5.84 (2.88)** | 2.27 (0.91) |

### 13C-glucose incubation experiment

To enable identification of substrate-induced active lytic virus populations, we traced ^13^C from added isotopically labelled glucose into isolated virome DNA (Supplementary Fig. S1). To accomplish this, two 100 g (dry-weight equivalent) subsamples of each of the eight soils were weighed into autoclaved centrifuge bottles and pre-incubated at 20°C for one week. We then added either 99 atom% ^13^C-glucose (Cambridge Isotope Labs catalog# CLM-1396-2) or a natural abundance ^12^C-glucose control to the microcosms at a rate of 0.5 mg glucose-C g dry soil^-1^. The glucose-C treatment solutions were added to the units such that the soils were brought to 65% of their water holding capacity. The solutions were added to field-moist soils already at ∼50% of their water holding capacity, which was intended to minimize the effects of added moisture on microbe-virus dynamics. A total of 16 units were incubated for the experiment (2 land uses × 2 isotope treatments × 4 replicate soils per land use = 16). The units were loosely capped to allow for gas exchange and were incubated in the dark at 20°C for one week. Because the added glucose likely propagated through primary and secondary consumers during the one-week incubation period, the results should be interpreted as integrated glucose responses over the experimental timeframe rather than initial microbe-virus interactions. Microbial respiratory responses to the glucose additions were determined using separate microcosm static incubations, using the same glucose-C addition rates plus an additional water-only control microcosm for each soil (29, 44). CO_2_ concentration in the headspace of the microcosms was measured using a Q-S151 infrared gas analyzer (Qubit Systems Inc. Kingston, ON, Canada).

### Virome DNA extraction and density gradient ultracentrifugation

After the incubation, we extracted virions from the soils using two successive extractions with 100 ml of amended potassium citrate solution (1% potassium citrate, 150 mM MgSO_4_, 13.7 mM NaCl, 0.27 mM KCl, 1 mM Na_2_HPO_4_, 0.18 mM KH_2_PO_4_) (14). For each successive extraction, solution was added directly to the experimental units, the units were shaken for 30 min on low speed on a reciprocal shaker and then centrifuged at 3500 × *g* for 20 min at 4°C to pellet soil particles. The two supernatants from each unit were pooled and microbial cells depleted by passing through 0.2 µm pore size polyethersulfone (PES) filters (VWR catalog# 10040-440). We concentrated virions in the filtered supernatants ∼500-fold to a final volume of ∼400 µl using 100 kDa Pierce^TM^ centrifugal concentration units (Thermo Fisher Scientific, Waltham, MA, USA). We depleted unencapsulated DNA in the concentrates by adding 20 units of RQ1 DNase enzyme (Promega corp, Madison, WI, USA), incubating for 30 min at 37°C, then adding 20 µl of stop solution to terminate the reaction. Then, DNA was extracted from the DNase-treated virion concentrates using a Qiagen PowerSoil Pro kit (Qiagen, Hilden, Germany). We modified the manufacturer’s extraction protocol by substituting the bead lysis step with a 15 min heat lysis step at 65°C (15). DNA extracts were quantified on a Qubit 4.0 fluorometer (Life Technologies, Carlsbad, CA, USA).

The virome DNA samples from the incubations (0.2 – 1 µg DNA) were subjected to isopycnic CsCl density gradient centrifugation (1.69 g ml^-1^ starting density) in 5.1 ml QuickSeal tubes in a Beckman Optima L-100 XP ultracentrifuge equipped with a 100-ti rotor at 162,000 × g for 72 h (Beckman Coulter Inc., Brea, CA, USA). The CsCl density gradients were then fractionated into 24 ∼200 µl fractions. The density of the fractions was measured using a Mettler-Toledo MyBrix handheld refractometer (Mettler-Toledo, Columbus, OH, USA). DNA was recovered from the fractions via precipitation with PEG 8000 (45) and pelleted DNA was washed with 70% ethanol, then resuspended in 40 µl of elution buffer (10 mM Tris-Cl, pH 8.5). DNA in the recovered fractions was quantified using a Qubit 4.0 fluorometer. The fractions were then pooled into three density ranges: high density DNA (> 1.71 g ml^-1^) (46, 47), medium density DNA (1.69 – 1.71 g ml^-1^) and low density DNA (1.65 – 1.69 g ml^-1^) (Fig. 1). This pooling step was done to ensure sufficient DNA was present for metagenome library prep. The pooling step also involved DNA size selection (>2,000 bp) using the Zymo DNA clean and concentrator kit (Zymo Research, Irvine, CA, USA) to further deplete fragmented microbial DNA in the samples. This resulted in 48 virome DNA samples total from the SIP fractions (2 land uses × 2 isotope treatments × 4 replicate soils× 3 density fractions = 48).

**Figure 1:**
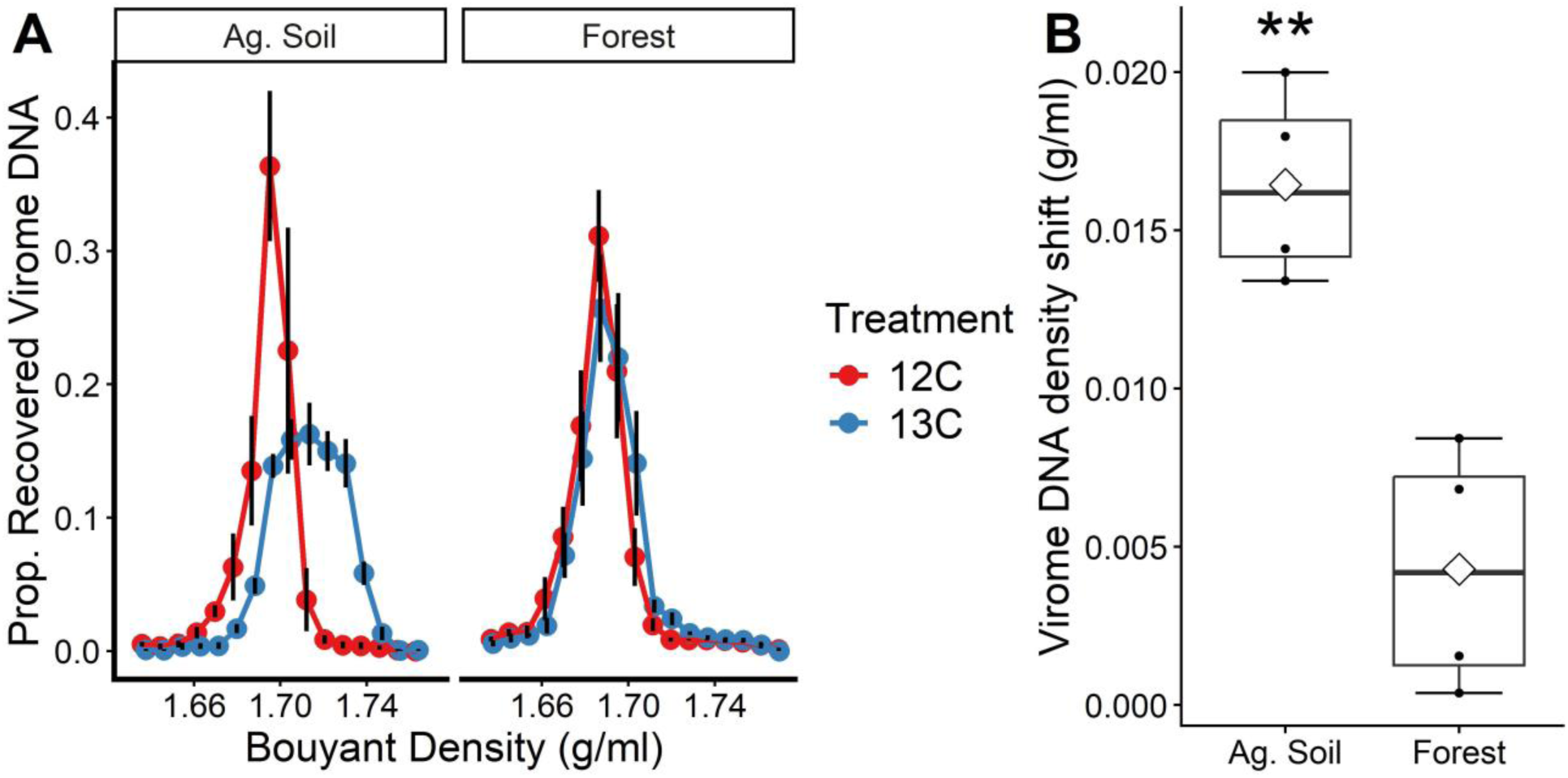
Virome DNA density shifts in the agricultural and forest soils. Panel (A) shows the virome DNA density curves averaged across the four replicates of each soil for each isotope treatment. The y-axis values were calculated as the proportion of the total virome DNA isolated from the microcosms that was recovered in each respective density fraction. Panel (B) shows the change in weighted average virome DNA density (^13^C treatment - ^12^C treatment) for each of the four replicate soils from each ecosystem (B). Error bars in (A) show one standard error of the mean. Diamonds in (B) indicate treatment means. Asterisks in (B) indicate statistically greater density shifts (one-way ANOVA) at the following significance level: ** p < 0.01.

### Sequencing and bioinformatics analyses

Virus metagenome (virome) libraries were prepared for sequencing using Nextera XT kits (Illumina, San Diego, CA, USA) and the libraries sequenced on the Illumina NovaSeqX platform with 2 × 151 bp reads at the US Department of Energy Joint Genome Institute (JGI). NCBI accession numbers for the raw sequences are provided in Supplementary Table S1. Initial processing of the raw sequence reads was done using the automated JGI QC pipeline for virus metagenomes. Briefly, this involved using BBDuk (https://jgi.doe.gov/data-and-tools/bbtools/) to remove optical duplicates, low quality bases, reads < 51bp, and reads aligning to common microbial and eukaryote contaminants. Based on the QC results, one pooled medium density fraction from the agricultural site was excluded from further analysis due to contamination detected. The quality-filtered reads were then assembled using metaSPAdes (48) (version 4.0). We supplemented the individual assemblies with two site-level co-assemblies performed using megahit (49) (version 1.2.9) in ‘meta-large’ mode. Sequence depth and assembly statistics for the samples are provided in Supplementary Table S2.

The contigs from all assemblies were combined and contigs >10,000 bp were passed to geNomad (50) (version 1.9.4) in ‘--composition virome’ mode for virus genome prediction and taxonomic annotation. The predicted virus genomes were checked for quality and completeness using CheckV (51) (version 1.0.3). Based on the CheckV results, we removed all proviral sequences and all low, medium, and undetermined quality virus contigs. We also removed genomes with no virus hallmark genes detected as well as genomes containing more than five host genes (41). All downstream analyses were thus performed on complete or high-quality (>90% complete) virus genomes. The high-quality genomes included 30,921 contigs out of the 294,881 putative viral contigs initially identified by geNomad. The resulting genome set was dereplicated using vClust (52) using the standard cutoff for virus genomes: 95% average nucleotide identify over 85% of the shortest sequence. This resulted in a total of 15,711 dereplicated genomes, *i.e.*, virus operational taxonomic units (vOTUs). We then aligned the quality-filtered sequence reads from each sample to the vOTU genome sequences using minimap2 (53) and calculated coverage and relative abundance statistics using coverM (54) (version 0.7.0), specifying a minimum covered fraction of 0.75. Host taxonomy of the virus genomes was determined using iPHoP (55) (version 1.3.3) using a minimum score of 75.

Following the recommendation of the developer, we only interpret host taxonomy at the family level when using a minimum score of 75. We constructed gene-sharing networks for the vOTUs within each soil type using vConTACT3 (56) (version 3.1.8). VIBRANT (57) (version 1.2.1) was used to categorize vOTUs as lytic or potentially lysogenic on the basis of lysogeny-associated genes in the genomes (*e.g.*, integrases, excisionases, transposases, recombinases, etc.). Putative auxiliary metabolism genes (AMGs) in the genomes were annotated using DRAM-v (version 1.5) (58). Coverage and sequence diversity of the virus metagenomes was determined using Nonpareil read redundancy analysis (version 3.5.5) (59).

### Statistical Analyses

We calculated ^13^C excess atom fraction (EAF) values of the vOTUs using the quantitative stable isotope probing (qSIP) equations (60) as implemented by the SIPmg R package (61).

SIPmg is optimized for qSIP analysis of genomes, taking into account the observed GC content of genomes while also adjusting bootstrapped EAF values for false discovery rates. Prior to qSIP analyses, we removed low-frequency vOTUs only observed in a single density fraction of a single replicate. For vOTU absolute abundance estimation, we used the ‘relative_coverage’ method recommended by the SIPmg developers (61). We then used SIPmg to estimate the EAF values and calculated 99% confidence intervals for the atom fraction excess values using 10,000 bootstrap comparisons. Virus genomes were considered to be isotopically labelled if the lower bound of the 99% confidence interval was above 0. We note here that our strategy of pooling density fractions reduces the resolution of the qSIP analyses (62), thus, our method may only confidently detect highly ^13^C-enriched vOTUs, whereas weakly labelled viruses may be missed.

To test for variation among microbial taxonomic groups at different taxonomic levels in terms of the ^13^C-EAF values of their associated vOTUs, we used generalized linear models (Gamma distribution, log-link function). To determine if there was microbial host phylogenetic organization in the ^13^C-EAF of vOTUs, we tested for phylogenetic signal using Blomberg’s *K* and Pagel’s λ statistics (63, 64). To analyze vOTU lytic activity in relation to host microbial growth traits, we calculated the average maximum growth rates of each host bacterial family using predicted minimum doubling times of bacterial genomes provided in the EGGO database (65, 66). To compare the gene content of ^13^C-enriched vOTUs and unenriched vOTU genomes, we used hypergeometric tests in the NEAT R package (67) to test whether vOTUs in the two enrichment categories had more or fewer links in gene-sharing networks than expected by chance. The networks were visualized using the igraph R package (68). To compare the sets of ^13^C enriched vs. unenriched vOTUs in terms of the prevalence of different genome characteristics (*i.e*., lysogeny genes, AMG categories), we used *z*-tests of proportions.

Correlations between variables were analyzed using Pearson correlation analysis. To compare the two soil types in terms of soil properties, virome DNA density shifts, Nonpareil sequence diversity, and respiratory responses to the glucose additions, we used one-way analysis of variance (ANOVA). All data and analysis code are available on figshare: https://doi.org/10.6084/m9.figshare.31416320.

## Results and Discussion

### Characterizing glucose-responsive active soil virus communities with virome DNA-SIP

To investigate the glucose-responsive active lytic virus communities in soils from contrasting environmental contexts, we collected soils from an undisturbed forest ecosystem and a highly disturbed long-term agricultural field. The soils varied dramatically in their physicochemical properties; the forest soils had more than 2-fold greater C content and C:N ratios while the agricultural soils had significantly higher pH and 58-fold greater extractable N (Table 1). We isolated virome DNA from the soils after incubating with either ^13^C-glucose or a natural abundance ^12^C-glucose control. We reasoned that since the C that comprises virions originates from host organisms, the detection of isotopically labelled virus genetic material is indicative of active lytic replication of virus populations in microbial hosts that incorporated the glucose. We observed increased virome DNA density in all the ^13^C-incubated soils, indicating successful isotopic labelling of virome DNA (Fig. 1). However, the degree of isotope labelling varied greatly between the two soils; the density shifts observed in the agricultural soils were ∼4-fold greater on average than those observed in the forest soils (Fig. 1A, B). This indicates greater viral lysis (thus greater virus DNA isotope labelling) in response to glucose-C additions in the agricultural samples compared to the forest soil samples.

To characterize the glucose-responsive active lytic virus communities in the two soils, we conducted metagenome sequencing of low, medium, and high-density virome DNA density fractions, followed by quantitative stable isotope probing (qSIP) analysis of the assembled virus genomes (60, 61). We recovered 15,711 high-quality (>90% complete), non-redundant virus genomes (vOTUs) from the virus metagenome sequence data. Of these vOTUs, ∼90% were unique to the forest soils. Only three vOTUs occurred in both soil types (Supplementary Fig. S2), despite the fact that 177 prokaryote ASVs (∼5% of ASVs) overlapped between the soils (Supplementary Fig. S3). This reflects the greater beta diversity in virus communities compared with prokaryote communities that has been previously observed in soil (32, 40). vOTU richness values averaged ∼430 vOTUs per sample in the agricultural soils and ∼2,800 vOTUs per sample in the forest soils (Supplementary Fig. S4). The much higher vOTU richness we observed in the forest soils was unexpected and is despite the similar levels of prokaryote richness we observed between the soil types (Supplementary Fig. S3). Analysis of sequence read redundancy (59) revealed that the agricultural soil virus metagenomes had higher coverage than the forest soil virus metagenomes (mostly >90% vs. ∼60-80%) but substantially lower sequence-level diversity than the forest soils (Supplementary Fig. S5, S6). This implies that lack of coverage did not limit our ability to identify vOTUs in the agricultural soils and supports our observation of greater vOTU diversity in the forest soils. One possible explanation for the difference in vOTU richness between the soils is that soil disturbance disproportionately affects viruses compared with microorganisms, thus leading to reduced vOTU alpha diversity in the agricultural soils.

After removing low-frequency vOTUs, we conducted qSIP analysis to estimate bootstrapped ^13^C excess atom fraction (EAF) values for the vOTUs (60, 61). This analysis revealed diverse, active (*i.e.*, ^13^C-incorporating) soil virus communities in both ecosystems, with 5,353 active populations identified in total out of the 11,058 vOTUs with bootstrapped EAF estimates (Fig. 2). However, the patterns of isotope labelling varied greatly between the two soil types. In line with the virome DNA density shifts, the agricultural soil vOTUs exhibited much greater ^13^C-EAF values, indicating greater virus lytic activity in response to glucose addition (Fig. 2A). In those soils, 77% of vOTUs with bootstrapped EAF values (754/973) were ^13^C-incorporators and the median EAF of vOTUs was 0.24. This is in contrast to the forest soils, where 46% of vOTUs (4,599/10,085) were ^13^C-incorporators and the median EAF of vOTUs was 0.02 (Fig. 2B). We suggest that these differences can be attributed to distinct microbial responses to the glucose additions between the soils: the agricultural soils had 2.6-fold greater respiratory responses to glucose (relative to no-glucose controls) compared to the forest soils (Table 1, Supplementary Fig. S7), which, in turn, likely promoted greater viral lysis of the glucose-responsive agricultural soil microorganisms. In turn, it is likely that the differences in the glucose respiratory responses are due to greater C-limitation in the agricultural soils compared to the C-rich forest soils (Table 1). However, the soils also varied in other aspects, *e.g*., pH, extractable N (Table 1), thus we cannot rule out the possibility that other factors influenced the differences in glucose-induced virus activity. Regardless, these results suggest fundamental differences in host-virus interactions among soil contexts, especially in soil microsites subject to frequent labile C additions, *e.g.*, the rhizosphere. Overall, our results point to a central role of C availability/inputs in mediating host-virus interactions in soil, similar to other recent studies (23, 29).

**Figure 2:**
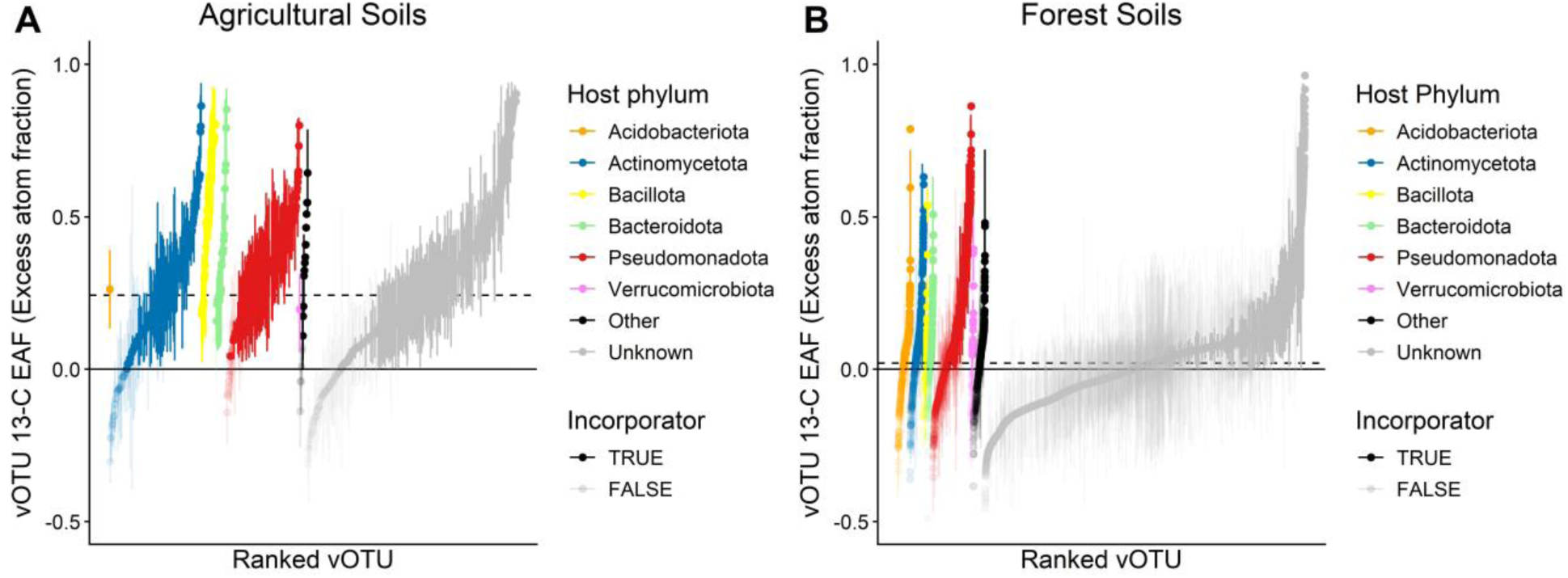
vOTU ^13^C excess atom fraction (EAF) in the agricultural and forest soils with 99% confidence intervals (error bars) calculated from 10,000 bootstrap comparisons. ^13^C-incorporating (*i.e.*, active lytic) vOTUs have 99% confidence intervals with the lower bound above the zero line. 77% of vOTUs in the agricultural soils were ^13^C incorporators (A) while 46% of vOTUs in the forest soils were ^13^C incorporators (B). The dashed lines indicate the median vOTU ^13^C EAF values for each soil type. The “Other” category represents low abundance microbial phyla with only one or few associated vOTUs identified.

### Characteristics of the microbial hosts of glucose-responsive active soil viruses

Microbial host prediction for the vOTUs revealed that the glucose-induced active viruses were linked to hosts from 197 prokaryote families across 28 phyla. However, it is notable that the majority of the active vOTUs had no predicted microbial host, particularly in the forest soils, where 75% had no host prediction (compared to ∼50% in the agricultural soils) (Fig. 2). Poor identification of microbial hosts is a known limitation in the field of soil virology (8) and the difference in annotation success between the soils likely reflects the greater relative abundance of more oligotrophic, poorly characterized microbial groups in the forest soils (*e.g.*, Acidobacteriota, Verrucomicrobiota lineages). In the agricultural soils, nearly all ^13^C-incorporating vOTUs with host taxonomy annotations infected hosts from only four prokaryotic phyla: Actinomycetota (153 vOTUs), Bacillota (34 vOTUs), Bacteroidota (25 vOTUs), and Pseudomonadota (155 vOTUs) (Fig. 2A). In the forest soils, the largest number of ^13^C-incorporating vOTUs with host taxonomic annotations had Pseudomonadota hosts (556 vOTUs), followed by Actinomycetota (202 vOTUs) and Acidobacteriota (153 vOTUs) (Fig. 2B). The differences in the host taxonomy of active lytic vOTUs between the two soils reflected differences in the underlying microbiomes of the soils, *e.g.*, the forest soils had greater representation of Acidobacteriota and Verrucomicrobiota both in their microbiomes and within the active vOTU hosts (Supplementary Fig. S8). Notable microbial host families with high vOTU ^13^C enrichments included the *Bacillaceae* (Bacillota), *Micrococcaceae* (Actinomycetota), *Bacteroidaceae* (Bacteroidota), and *Vibrionaceae* (Pseudomonadota).

To evaluate phylogenetic patterns of glucose-induced virus lytic activity among host microbial groups, we pruned the genome taxonomy database (GTDB) tree (69) to only include microbial taxa present in our study and tested for host phylogenetic signal in vOTU ^13^C-EAF (63, 64) (Fig. 3). This analysis revealed significant microbial phylogenetic signal in virus lytic activity only in the agricultural soil communities (Fig. 3). This indicates that there is measurable variation in the degree of viral lysis among microbial phylogenetic groups in the agricultural soils. The lack of evident phylogenetic organization in the forest soils likely relates to the overall much lower virus activity present in those soils under the experimental conditions in this study. Regardless, these results demonstrate that both microbial evolutionary history and soil environmental context influence the degree of viral lysis following resource inputs to soil. To further explore patterns in virus lytic activity among microbial taxa, we examined within- and among-group variation in vOTU ^13^C-EAF among host microbial taxonomic groups at different taxonomic levels. Similar to prior work on microbial ^13^C-glucose incorporation (70), microbial host taxonomy at all taxonomic levels explained significant amounts of variation in vOTU ^13^C EAF values (Fig. 4) (Supplementary Table S3). However, microbial taxonomy explained more variation in vOTU ^13^C-EAF in the agricultural soils than in the forest soils (Supplementary Table S3), which is consistent with the differences in phylogenetic signal of vOTU ^13^C-EAF between the soils (Fig. 3). Further, in both soils, finer microbial taxonomic levels explained greater proportions of variation in vOTU ^13^C-EAF than broader taxonomic levels (up to 40% and 32% at the family level for the agricultural and forest soil, respectively) (Supplementary Table S3), which is also similar to prior microbial SIP studies (70–72). In general, viruses of Bacillota and Bacteroidota hosts exhibited the greatest ^13^C-EAF in the agricultural soils, while in the forest soils, viruses of Pseudomonadota hosts exhibited the greatest ^13^C-EAF (Fig. 4A, B).

**Figure 3:**
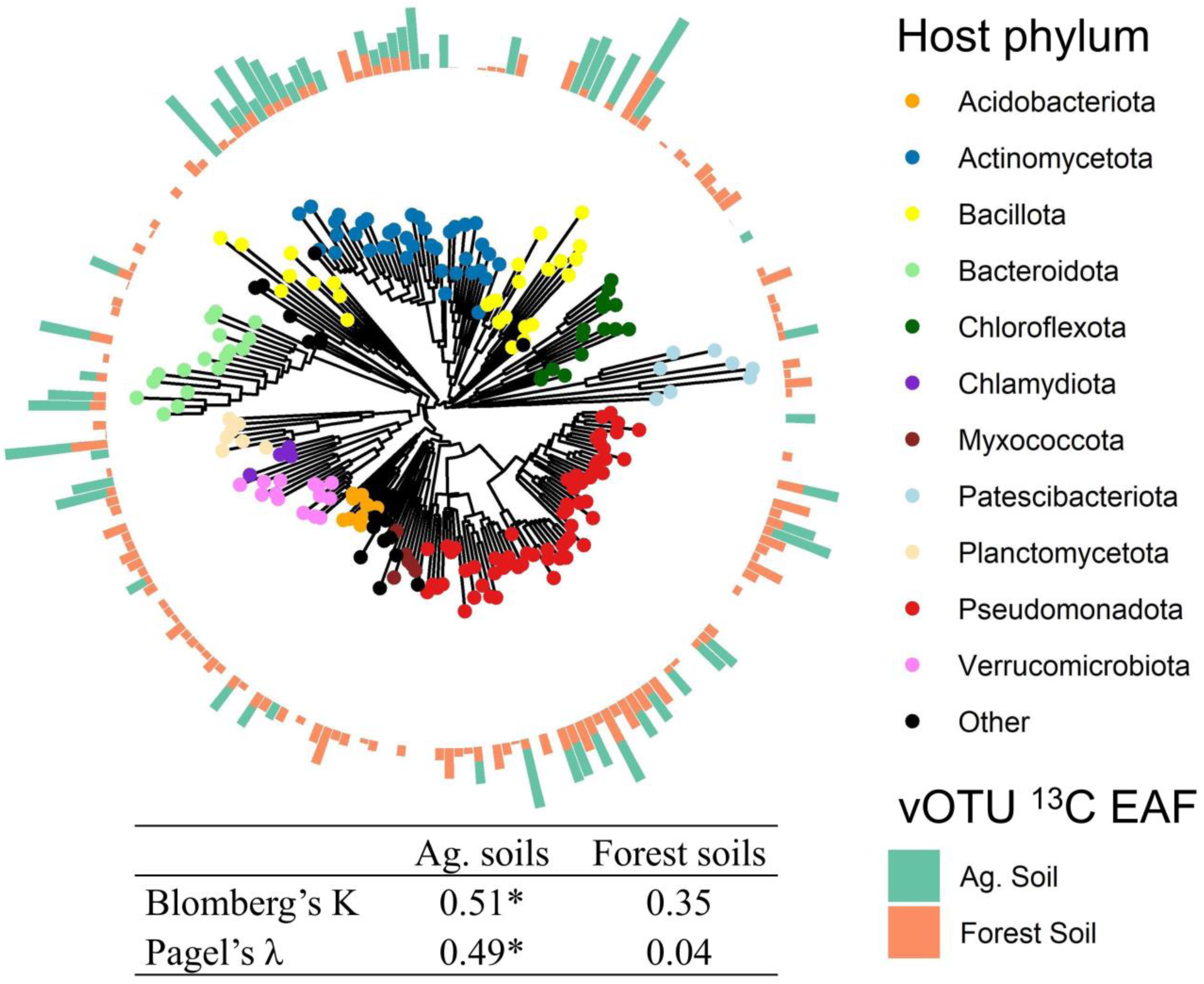
Phylogenetic patterns in lytic virus activity (vOTU ^13^C-EAF) among microbial host taxa. The genome taxonomy database (GTDB) tree was pruned to only include taxa present in our study. Bars around the tree are proportional to the ^13^C-EAF values of vOTUs linked to the microbial host families in the agricultural soils (green bars) and forest soils (orange bars). Host phylogenetic signal in vOTU ^13^C EAF was analyzed using Blomberg’s K and Pagel’s λ statistics. Asterisks indicate statistically significant phylogenetic signal detected (*p* < 0.05).

**Figure 4:**
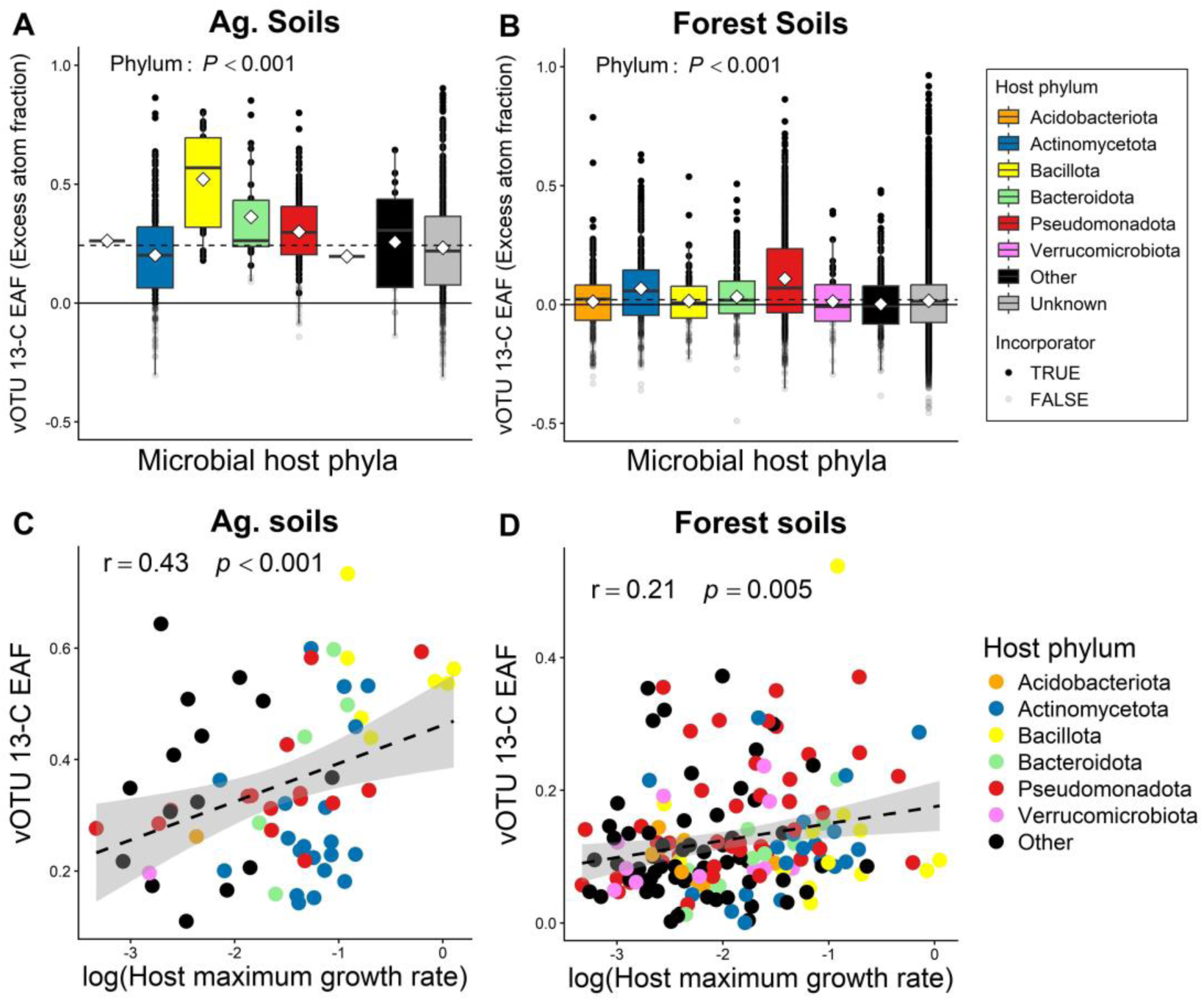
Variation in vOTU ^13^C-EAF among microbial taxonomic and life history groups. Shown are the distributions of vOTU ^13^C EAF values among microbial host phyla of the active vOTUs in the agricultural (A) and forest (B) soils. Diamond symbols on (A) and (B) indicate the mean vOTU ^13^C EAF for each host phylum while the dashed lines show the median value for each soil type. *P* values on (A) and (B) are from generalized linear models. Also shown are the vOTU ^13^C EAF values in comparison to the maximum growth rate of the microbial host for the agricultural (C) and forest (D) soils. For panels (C) and (D) both vOTU ^13^C EAF and host growth rates are aggregated at the host family level, with 64 families represented in the agricultural soil and 169 families represented in the forest soil. Host maximum growth rates were derived from the EGGO database of predicted minimum doubling times for microbial genomes. Correlation coefficients and *p* values on (C) and (D) are from Pearson correlation analysis.

These results relate to our original hypothesis that the degree of glucose-induced viral lysis would be greatest in highly copiotrophic taxa. Indeed, the microbial host phyla of the most highly ^13^C-enriched vOTUs (*e.g.*, Bacillota, Pseudomonadota) have been identified by several previous studies as having many copiotrophic members (65, 71, 73). To further explore this, we compared the vOTU ^13^C-EAF values to the maximum growth rate of each microbial family, calculated based on the minimum doubling times of microbial genomes provided in the EGGO database (66). We observed significant, positive correlations between microbial host maximum growth rate and vOTU ^13^C-EAF in both soils (Fig. 4 C, D), though the correlation was weaker in the forest ecosystem (Fig. 4D). This is consistent with the weaker influence of host microbial taxonomy/phylogeny in explaining vOTU ^13^C-EAF in the forest soils. This could be because the low nutrient availability (high C:N) forest soils constrained both actual microbial growth in response to glucose as well as maximum microbial growth rates, which were 44% lower on average in the forest soil host taxa than the agricultural hosts. In any case, these results provide support for our hypothesis regarding the links between microbial traits (growth potential) and virus lytic activity in the context of labile substrate additions and potentially explain the overall greater virus activity that we observed in the agricultural soils compared to the forest soils.

Though the correlations between microbial growth potential and vOTU activity were modest in strength, our ability to detect the relationships at all is notable given the inherently noisy nature of the quantitative variables involved, *e.g*., the aforementioned limited resolution of our ^13^C-EAF values along with the uncertain relationships between microbial growth potential and actual growth in our experimental units. Further, the association between microbial copiotrophy and viral lysis is consistent with the “cull-the-winner” model of virus replication, though this result should be evaluated cautiously since there are multiple alternative scenarios in which this correlation could manifest. For example, this relationship could simply reflect a situation in which viral production is proportional to microbial host population growth in response to the glucose additions, as opposed to the disproportional lysis of microbial copiotrophs and associated density-dependent dynamics that are predicted by “cull-the-winner” (32). Disentangling these possibilities would require observation of temporal microbial and viral population changes, whereas our study only provides a single end-point snapshot of host-virus interactions. Thus, while our results support the specific hypotheses we initially posed, we are unable to provide strong support for the broader hypotheses regarding viral replication strategies in soil.

An additional caveat is that since our isotope labelling results are substrate-dependent, the relationships demonstrated here are only applicable to limited subsets of glucose-responsive microbial copiotrophs and their associated viruses. Thus, while it appears that the degree of microbial copiotrophy is related to the degree of lytic virus activity within glucose-responsive taxa, we cannot draw conclusions regarding links between microbial growth potential and viral lysis in the soil microbiome broadly. Regardless, the relationships we observed among soil context, microbial growth traits, and viral activity suggest that factors that deplete soil C stocks and/or increase N availability (*e.g*., land use change, climate change, etc.) will increase the prevalence of lytic virus replication by promoting the growth of copiotrophic microbial hosts with boom-bust adapted life strategies following labile C additions (74). Furthermore, such increases in lytic replication could accelerate further soil C losses through microbial lysate-induced priming of organic matter decomposition (29).

### Characteristics of the active lytic virus genomes

Nearly all of the vOTUs in our dataset were identified as double-stranded DNA phages in virus class Caudoviricetes (>99%), though several other virus classes were present in very low relative abundances, including Maveriviricites, Megaviricetes, Quintoviricetes, and Tectiliviricetes (Supplementary Fig. S9). Very few of the vOTUs had annotations at lower taxonomic levels, *e.g.*, only ∼3% of vOTUs in each sample were assigned to virus families on average. The most common virus families identified were Zobellviridae (1.8% of vOTUs) and Autographiviridae (0.8% of vOTUs), both in class Caudoviricetes. These limitations reflect the well-documented challenges of classifying virus genomes assembled from environmental samples (75). Though vOTU taxonomic information was limited, we did observe evidence of distinct genome characteristics of the ^13^C-enriched vOTU genomes. In both soils, hypergeometric tests showed that ^13^C-enriched vOTUs had fewer links with unenriched vOTUs in gene-sharing networks than expected by chance (both *p* < 0.001, Supplementary Fig. S10).

This suggests that there may be unique functional characteristics of the active vOTU genomes. To explore this further, we conducted functional annotation of the vOTU genomes and compared the active, ^13^C-enriched genomes to the unenriched vOTU genomes. We reiterate here that our categorization of vOTUs as ^13^C-enriched vs. unenriched is imperfect due to the limited resolution of our qSIP analysis. Nonetheless, comparing vOTUs coarsely distinguished as more vs. less active is useful for revealing the genomic features of viruses with high substrate-induced lytic activity.

We first identified potentially lysogenic vOTUs based on the presence of lysogeny-associated genes in the genomes (57), which revealed that only 23% of the vOTUs in our dataset overall were potentially lysogenic. This is consistent with prior work showing that virions isolated from soil are mostly identified as strictly lytic (17, 24, 32). Interestingly, the proportion of potentially lysogenic vOTUs was much greater in the agricultural soils (52%) than in the forest soils (20%) (Fig. 5A), despite the greater lytic activity in response to glucose addition we observed in the agricultural soils. Furthermore, *z*-tests revealed that a significantly greater proportion of the active ^13^C-enriched vOTUs were potentially lysogenic compared to the unenriched vOTUs: 53% vs. 46% and 25% vs. 16% in the agricultural and forest soils, respectively (Fig. 5A). That said, statistical power was limited for the agricultural soil analysis due to a much smaller number of vOTUs present (especially in the non-enriched category).

**Figure 5:**
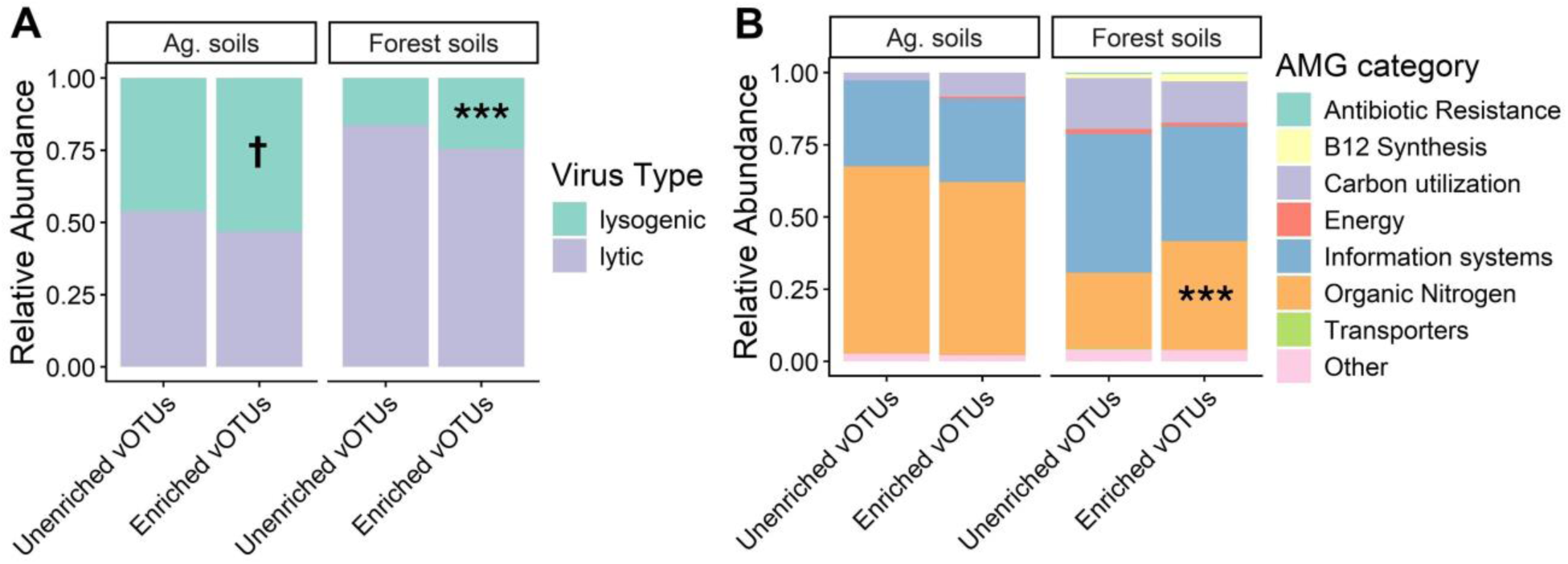
Comparisons of the active ^13^C enriched and unenriched vOTU genome characteristics from the two soil types. Panel (A) shows the relative abundance of potentially lysogenic viruses within the sets of enriched (active) and unenriched vOTUs across both soil types on the basis of lysogeny-associated genes detected in the genomes (integrases, excisionases, recombinases, transposases, etc.). Panel (B) indicates the relative abundances of putative AMG categories (annotated with DRAM-v) within the sets of enriched and unenriched vOTUs across both soil types. Symbols indicate significantly higher proportions (*z*-tests) at the following levels: † *p* < 0.1, \*\*\**p* < 0.001

While intriguing, it is worth noting that these analyses are limited by the fact that our bioinformatics approach likely does not perfectly distinguish between potentially lysogenic vs. strictly lytic vOTUs since there are many alternative modes of temperate infection that do not require canonical lysogeny genes (*e.g.*, integrases) (76). Nonetheless, these patterns suggest that virus communities with high substrate-induced lytic potential can also have high potential for lysogeny. Overall, our results suggest that induction of lysogenic viruses to lysis by labile C additions may have important effects on soil microbiomes and their biogeochemical functions, consistent with prior work that has demonstrated high lytic potential and large burst sizes in some communities of lysogenic soil viruses (77, 78).

To further explore the functional characteristics of the active vOTU genomes, we annotated potential auxiliary metabolism genes (AMGs) in the genomes. Because our strict CheckV filtering of viral contigs removed the great majority of host contamination from the vOTU genomes prior to AMG annotation (see Methods), we did not perform further manual curation of the AMGs identified. Nonetheless, it is important to note here that AMG annotation of virus genomes can suffer from high rates of false positives (79), thus the AMGs described here should be considered as putative in nature. DRAM-v (58) identified 2,013 putative AMGs across 1,782 vOTUs in our dataset. The majority of the identified putative AMGs had functions in cellular information systems (*e.g*., nucleotide biosynthesis), organic N acquisition (*e.g.*, protein and amino acid degradation) and C acquisition (*e.g*., CAZymes) (Fig. 5B). Though only 10-16% of genomes within each enrichment category × soil type combination had putative AMGs, we did observe differences between the ^13^C-enriched vs. non-enriched vOTU genomes in the prevalences of some notable AMG categories. Specifically, AMGs in the ^13^C-enriched vOTUs from the agricultural soils had greater representation of C-acquisition genes while enriched vOTU genomes from the forest soil had greater prevalence N-acquisition genes (Fig. 5B). These patterns achieved statistical significance in the forest soils (Fig. 5B), whereas robust statistical analysis in the agricultural soils was precluded by the fact that few unenriched vOTUs in those soils contained putative AMGs. In any case, the differences in putative AMG relative abundances we observed in the active vOTUs appear to reflect the element limitations of the two soils, *i.e.*, the active lytic viruses may also have enhanced potential to alleviate microbial C- and N-limitation in the agricultural and forest soils, respectively. While preliminary in nature, these patterns suggest that substrate-induced lytic replication may not come at the cost of promoting host fitness and survival through lysogenic incorporation of beneficial AMGs into host genomes. Thus, it is possible that highly active lytic virus populations that “cull-the-winner” may also initially “fuel-the-winner” through AMG incorporation into hosts prior to being induced to lysis (80).

## Conclusions

By tracing ^13^C-glucose from host microorganisms into virus genomic DNA, our study provides new insights into microbe-virus interactions in soil. First, our study demonstrates that substrate-induced virus activity can vary greatly depending on soil environmental context.

Related to this, it is important to reiterate that our study investigated virus lytic activity following glucose addition, which explains the different responses observed between the C-limited agricultural soils and the high C:N forest soils in our study. On that basis, one might expect that an alternative virome DNA SIP experiment using ^15^N additions would reveal opposite patterns in virus DNA isotope labelling between soil types than we observed here. Nevertheless, these results suggest distinct patterns of viral lysis among ecosystem types in soil habitats that receive frequent labile C additions, *e.g.*, the rhizosphere. An alternative isotope labelling approach would be to use H_218_O to identify substrate-independent active virus populations, as was done in one previous study (41). That study was broadly consistent with ours in that both revealed active viruses to have primarily Actinomycetota and Pseudomonadota hosts, though our study revealed far more active virus populations. This suggests that addition of labile ^13^C compounds may have greater potential to reveal diverse communities of lytic viruses, though with the additional caveats inherent to substrate addition experiments. Nonetheless, the two studies together demonstrate the feasibility and benefits of combining DNA-SIP with viromics.

Another intriguing finding of our study is the link we observed between microbial growth potential and the degree of virus lytic activity. This observation is consistent with the prevailing hypotheses of virus replication in soil (*e.g.*, “cull-the-winner”), though additional temporal studies of microbe/virus population dynamics are required to provide strong support.

Unexpectedly, we also observed that the viruses that lysed hosts following glucose addition were more likely to be potentially lysogenic. It is possible that many of the active viruses we detected had been lysogenic prior to being induced to lysis by the glucose additions, a process that was potentially also mediated by quorum-sensing in the growing prokaryote populations (81). In addition, putative AMG profiles in the vOTU genomes suggested that substrate-induced lytic virus populations may initially benefit host fitness through AMG incorporation into host genomes, which should be investigated in future work. An additional important area of future research is to extend our approach to investigate ssDNA and RNA virus activity in soil (6, 82–84). In any case, our study is an important contribution to soil virology, identifying substrate-responsive active lytic viruses and linking their activity to characteristics of their microbial hosts. In addition, our study demonstrates a powerful framework that can be extended to elucidate microbe-virus interactions in soil and their contributions to biogeochemical processes across terrestrial ecosystem contexts.

## Acknowledgements

The work (proposal: 511094, award DOI: 10.46936/10.25585/60013052) conducted by the U.S. Department of Energy Joint Genome Institute (https://ror.org/04xm1d337), a DOE Office of Science User Facility, is supported by the Office of Science of the U.S. Department of Energy operated under Contract No. DE-AC02-05CH11231. We thank Dr. Steve McBride for his helpful comments on an early draft of this manuscript.

## Data Availability Statement

NCBI Accession numbers for the raw virus metagenome sequence data are provided in Supplementary Table S1. All other data and analysis scripts are available on figshare: https://doi.org/10.6084/m9.figshare.31416320

## References

1. Kuzyakov Y, Mason-Jones K. 2018. Viruses in soil: Nano-scale undead drivers of microbial life, biogeochemical turnover and ecosystem functions. Soil Biology and Biochemistry 127:305–317.

2. Cobián Güemes AG, Youle M, Cantú VA, Felts B, Nulton J, Rohwer F. 2016. Viruses as Winners in the Game of Life. Annual Review of Virology 3:197–214.

3. Quirós P, Sala-Comorera L, Gómez-Gómez C, Ramos-Barbero MD, Rodríguez-Rubio L, Vique G, Yance-Chávez T, Atarés S, García-Gutierrez S, García-Marco S, Vallejo A, Salaet I, Muniesa M. 2023. Identification of a virulent phage infecting species of Nitrosomonas. 5. ISME J 17:645–648.

4. Barnett SE, Buckley DH. 2023. Metagenomic stable isotope probing reveals bacteriophage participation in soil carbon cycling. Environmental Microbiology 25:1785–1795.

5. Weinbauer MG, Rassoulzadegan F. 2004. Are viruses driving microbial diversification and diversity? Environmental Microbiology 6:1–11.

6. Williamson KE, Fuhrmann JJ, Wommack KE, Radosevich M. 2017. Viruses in Soil Ecosystems: An Unknown Quantity Within an Unexplored Territory. Annual Review of Virology 4:201–219.

7. Zimmerman AE, Graham EB, McDermott J, Hofmockel KS. 2024. Estimating the Importance of Viral Contributions to Soil Carbon Dynamics. Global Change Biology 30:e17524.

8. Hazard C, Anantharaman K, Hillary LS, Neri U, Roux S, Trubl G, Williamson K, Pett-Ridge J, Nicol GW, Emerson JB. 2025. Beneath the surface: Unsolved questions in soil virus ecology. Soil Biology and Biochemistry 205:109780.

9. Carreira C, Lønborg C, Acharya B, Aryal L, Buivydaite Z, Borim Corrêa F, Chen T, Lorenzen Elberg C, Emerson JB, Hillary L, Khadka RB, Langlois V, Mason-Jones K, Netherway T, Sutela S, Trubl G, wa Kang’eri A, Wang R, White RA, Winding A, Zhao T, Sapkota R. 2024. Integrating viruses into soil food web biogeochemistry. Nat Microbiol 1–11.

10. Suttle CA. 2007. Marine viruses — major players in the global ecosystem. Nat Rev Microbiol 5:801–812.

11. Fuhrman JA. 1999. Marine viruses and their biogeochemical and ecological effects. 6736. Nature 399:541–548.

12. Bowatte S, Newton PCD, Takahashi R, Kimura M. 2010. High frequency of virus-infected bacterial cells in a sheep grazed pasture soil in New Zealand. Soil Biology and Biochemistry 42:708–712.

13. Takahashi R, Bowatte S, Taki K, Ohashi Y, Asakawa S, Kimura M. 2011. High frequency of phage-infected bacterial cells in a rice field soil in Japan. Soil Science and Plant Nutrition 57:35–39.

14. Trubl G, Solonenko N, Chittick L, Solonenko SA, Rich VI, Sullivan MB. 2016. Optimization of viral resuspension methods for carbon-rich soils along a permafrost thaw gradient. PeerJ 4:e1999.

15. Trubl G, Roux S, Solonenko N, Li Y-F, Bolduc B, Rodríguez-Ramos J, Eloe-Fadrosh EA, Rich VI, Sullivan MB. 2019. Towards optimized viral metagenomes for double-stranded and single-stranded DNA viruses from challenging soils. PeerJ 7:e7265.

16. Göller PC, Haro-Moreno JM, Rodriguez-Valera F, Loessner MJ, Gómez-Sanz E. 2020. Uncovering a hidden diversity: optimized protocols for the extraction of dsDNA bacteriophages from soil. Microbiome 8:17.

17. Emerson JB, Roux S, Brum JR, Bolduc B, Woodcroft BJ, Jang HB, Singleton CM, Solden LM, Naas AE, Boyd JA, Hodgkins SB, Wilson RM, Trubl G, Li C, Frolking S, Pope PB, Wrighton KC, Crill PM, Chanton JP, Saleska SR, Tyson GW, Rich VI, Sullivan MB. 2018. Host-linked soil viral ecology along a permafrost thaw gradient. Nat Microbiol 3:870–880.

18. Emerson JB. 2019. Soil Viruses: A New Hope. mSystems 4:e00120–19.

19. Graham EB, Camargo AP, Wu R, Neches RY, Nolan M, Paez-Espino D, Kyrpides NC, Jansson JK, McDermott JE, Hofmockel KS. 2024. A global atlas of soil viruses reveals unexplored biodiversity and potential biogeochemical impacts. Nat Microbiol 9:1873–1883.

20. Ma B, Wang Y, Zhao K, Stirling E, Lv X, Yu Y, Hu L, Tang C, Wu C, Dong B, Xue R, Dahlgren RA, Tan X, Dai H, Zhu Y-G, Chu H, Xu J. 2024. Biogeographic patterns and drivers of soil viromes. Nat Ecol Evol 8:717–728.

21. Duan N, Li L, Liang X, McDearis R, Fine AK, Cheng Z, Zhuang J, Radosevich M, Schaeffer SM. 2022. Composition of soil viral and bacterial communities after long-term tillage, fertilization, and cover cropping management. Applied Soil Ecology 177:104510.

22. Duan N, Radosevich M, Zhuang J, DeBruyn JM, Staton M, Schaeffer SM. 2022. Identification of Novel Viruses and Their Microbial Hosts from Soils with Long-Term Nitrogen Fertilization and Cover Cropping Management. mSystems 7:e00571–22.

23. Wang S, López Arcondo JL, Xie N, Wang Y, Zhang Y, Radosevich M, Dutilh BE, Liang X. 2025. Exogenous carbon-to-nitrogen imbalance drives soil viral roles in microbial carbon mineralization and necromass accrual. Soil Biology and Biochemistry 210:109952.

24. Trubl G, Jang HB, Roux S, Emerson JB, Solonenko N, Vik DR, Solden L, Ellenbogen J, Runyon AT, Bolduc B, Woodcroft BJ, Saleska SR, Tyson GW, Wrighton KC, Sullivan MB, Rich VI. 2018. Soil Viruses Are Underexplored Players in Ecosystem Carbon Processing. mSystems 3.

25. Braga LPP, Spor A, Kot W, Breuil M-C, Hansen LH, Setubal JC, Philippot L. 2020. Impact of phages on soil bacterial communities and nitrogen availability under different assembly scenarios. Microbiome 8:52.

26. Albright MBN, Gallegos-Graves LV, Feeser KL, Montoya K, Emerson JB, Shakya M, Dunbar J. 2022. Experimental evidence for the impact of soil viruses on carbon cycling during surface plant litter decomposition. 1. ISME COMMUN 2:1–8.

27. Tong D, Wang Y, Yu H, Shen H, Dahlgren RA, Xu J. 2023. Viral lysing can alleviate microbial nutrient limitations and accumulate recalcitrant dissolved organic matter components in soil. ISME J 1–10.

28. Liang X, Sun S, Zhong Y, Zhang Y, Wang S, Wang Y, Xie N, Yang L, Radosevich M. 2024. Soil viruses reduce greenhouse gas emissions and promote microbial necromass accrual. bioRxiv 10.1101/2024.03.13.584929.

29. Osburn ED, Baer SG, Evans SE, McBride SG, Strickland MS. 2024. Effects of experimentally elevated virus abundance on soil carbon cycling across varying ecosystem types. Soil Biology and Biochemistry 198:109556.

30. Zhou Z, Liang X, Zhang N, Xie N, Huang Y, Zhou Y, Li B. 2024. The impact of soil viruses on organic carbon mineralization and microbial biomass turnover. Applied Soil Ecology 202:105554.

31. Castledine M, Buckling A. 2024. Critically evaluating the relative importance of phage in shaping microbial community composition. Trends in Microbiology 32:957–969.

32. Santos-Medellín C, Blazewicz SJ, Pett-Ridge J, Firestone MK, Emerson JB. 2023. Viral but not bacterial community successional patterns reflect extreme turnover shortly after rewetting dry soils. 11. Nat Ecol Evol 7:1809–1822.

33. Silveira CB, Rohwer FL. 2016. Piggyback-the-Winner in host-associated microbial communities. npj Biofilms Microbiomes 2:16010.

34. Hu C, Chen X, Wei W, Wallace D, Liu J, Zhang Y, Zhang L, Xu D, Batt J, Xiao X, Shi Q, Zheng Q, Ma R, Luo T, Jiao N, Zhang R. 2025. To kill or to piggyback: Switching of viral lysis-lysogeny strategies depending on host dynamics. Science of The Total Environment 959:178233.

35. Giovannoni S, Temperton B, Zhao Y. 2013. Giovannoni et al. reply. Nature 499:E4–E5.

36. Neufeld JD, Vohra J, Dumont MG, Lueders T, Manefield M, Friedrich MW, Murrell JC. 2007. DNA stable-isotope probing. 4. Nat Protoc 2:860–866.

37. Starr EP, Shi S, Blazewicz SJ, Koch BJ, Probst AJ, Hungate BA, Pett-Ridge J, Firestone MK, Banfield JF. 2021. Stable-Isotope-Informed, Genome-Resolved Metagenomics Uncovers Potential Cross-Kingdom Interactions in Rhizosphere Soil. mSphere 6:e00085–21.

38. Lee S, Sieradzki ET, Nicol GW, Hazard C. 2023. Propagation of viral genomes by replicating ammonia-oxidising archaea during soil nitrification. 2. ISME J 17:309–314.

39. Greenlon A, Sieradzki E, Zablocki O, Koch BJ, Foley MM, Kimbrel JA, Hungate BA, Blazewicz SJ, Nuccio EE, Sun CL, Chew A, Mancilla C-J, Sullivan MB, Firestone M, Pett-Ridge J, Banfield JF. 2022. Quantitative Stable-Isotope Probing (qSIP) with Metagenomics Links Microbial Physiology and Activity to Soil Moisture in Mediterranean-Climate Grassland Ecosystems. mSystems 7:e00417–22.

40. Santos-Medellin C, Zinke LA, ter Horst AM, Gelardi DL, Parikh SJ, Emerson JB. 2021. Viromes outperform total metagenomes in revealing the spatiotemporal patterns of agricultural soil viral communities. 7. ISME J 15:1956–1970.

41. Trubl G, Roux S, Kellom M, Vyshenska D, Tomatsu A, Singh K, Kimbrel JA, Eloe-Fadrosh E, Malmstrom RR, Pett-Ridge J, Blazewicz SJ. 2025. Tracking Persistence and Dynamics of Active Soil Viruses with SIP-Viromics. bioRxiv 10.1101/2025.05.25.655894.

42. Blevins RL, Thomas GW, Cornelius PL. 1977. Influence of No-tillage and Nitrogen Fertilization on Certain Soil Properties after 5 Years of Continuous Corn1. Agronomy Journal 69:383–386.

43. Sena KL, Williamson TN, Barton CD. 2021. The Robinson Forest environmental monitoring network: Long-term evaluation of streamflow and precipitation quantity and stream-water and bulk deposition chemistry in eastern Kentucky watersheds. Hydrological Processes 35:e14133.

44. West AW, Sparling GP. 1986. Modifications to the substrate-induced respiration method to permit measurement of microbial biomass in soils of differing water contents. Journal of Microbiological Methods 5:177–189.

45. Dunford EA, Neufeld JD. 2010. DNA Stable-Isotope Probing (DNA-SIP). J Vis Exp 2027.

46. Pepe-Ranney C, Campbell AN, Koechli CN, Berthrong S, Buckley DH. 2016. Unearthing the Ecology of Soil Microorganisms Using a High Resolution DNA-SIP Approach to Explore Cellulose and Xylose Metabolism in Soil. Front Microbiol 7.

47. Youngblut ND, Barnett SE, Buckley DH. 2018. SIPSim: A Modeling Toolkit to Predict Accuracy and Aid Design of DNA-SIP Experiments. Front Microbiol 9.

48. Nurk S, Meleshko D, Korobeynikov A, Pevzner PA. 2017. metaSPAdes: a new versatile metagenomic assembler. Genome Res 27:824–834.

49. Li D, Liu C-M, Luo R, Sadakane K, Lam T-W. 2015. MEGAHIT: an ultra-fast single-node solution for large and complex metagenomics assembly via succinct de Bruijn graph. Bioinformatics 31:1674–1676.

50. Camargo AP, Roux S, Schulz F, Babinski M, Xu Y, Hu B, Chain PSG, Nayfach S, Kyrpides NC. 2024. Identification of mobile genetic elements with geNomad. Nat Biotechnol 42:1303–1312.

51. Nayfach S, Camargo AP, Schulz F, Eloe-Fadrosh E, Roux S, Kyrpides NC. 2021. CheckV assesses the quality and completeness of metagenome-assembled viral genomes. Nat Biotechnol 39:578–585.

52. Zielezinski A, Gudyś A, Barylski J, Siminski K, Rozwalak P, Dutilh BE, Deorowicz S. 2025. Ultrafast and accurate sequence alignment and clustering of viral genomes. Nat Methods 22:1191–1194.

53. Li H. 2018. Minimap2: pairwise alignment for nucleotide sequences. Bioinformatics 34:3094–3100.

54. Aroney STN, Newell RJP, Nissen JN, Camargo AP, Tyson GW, Woodcroft BJ. 2025. CoverM: read alignment statistics for metagenomics. Bioinformatics 41:btaf147.

55. Roux S, Camargo AP, Coutinho FH, Dabdoub SM, Dutilh BE, Nayfach S, Tritt A. 2023. iPHoP: An integrated machine learning framework to maximize host prediction for metagenome-derived viruses of archaea and bacteria. PLOS Biology 21:e3002083.

56. Bolduc B, Zablocki O, Turner D, Jang HB, Guo J, Adriaenssens EM, Dutilh BE, Sullivan MB. 2025. Scalable and systematic hierarchical virus taxonomy with vConTACT3. bioRxiv 10.1101/2025.11.06.686974.

57. Kieft K, Zhou Z, Anantharaman K. 2020. VIBRANT: automated recovery, annotation and curation of microbial viruses, and evaluation of viral community function from genomic sequences. Microbiome 8:90.

58. Shaffer M, Borton MA, McGivern BB, Zayed AA, La Rosa SL, Solden LM, Liu P, Narrowe AB, Rodríguez-Ramos J, Bolduc B, Gazitúa MC, Daly RA, Smith GJ, Vik DR, Pope PB, Sullivan MB, Roux S, Wrighton KC. 2020. DRAM for distilling microbial metabolism to automate the curation of microbiome function. Nucleic Acids Res 48:8883–8900.

59. Rodriguez-R LM, Gunturu S, Tiedje JM, Cole JR, Konstantinidis KT. 2018. Nonpareil 3: Fast Estimation of Metagenomic Coverage and Sequence Diversity. mSystems 3:10.1128/msystems.00039-18.

60. Hungate BA, Mau RL, Schwartz E, Caporaso JG, Dijkstra P, van Gestel N, Koch BJ, Liu CM, McHugh TA, Marks JC, Morrissey EM, Price LB. 2015. Quantitative Microbial Ecology through Stable Isotope Probing. Appl Environ Microbiol 81:7570–7581.

61. Vyshenska D, Sampara P, Singh K, Tomatsu A, Kauffman WB, Nuccio EE, Blazewicz SJ, Pett-Ridge J, Louie KB, Varghese N, Kellom M, Clum A, Riley R, Roux S, Eloe-Fadrosh EA, Ziels RM, Malmstrom RR. 2023. A standardized quantitative analysis strategy for stable isotope probing metagenomics. mSystems 8:e01280–22.

62. Sieradzki ET, Koch BJ, Greenlon A, Sachdeva R, Malmstrom RR, Mau RL, Blazewicz SJ, Firestone MK, Hofmockel KS, Schwartz E, Hungate BA, Pett-Ridge J. 2020. Measurement Error and Resolution in Quantitative Stable Isotope Probing: Implications for Experimental Design. mSystems 5:10.1128/msystems.00151-20.

63. Blomberg SP, Garland T, Ives AR. 2003. Testing for phylogenetic signal in comparative data: behavioral traits are more labile. Evolution 57:717–745.

64. Pagel M. 1999. Inferring the historical patterns of biological evolution. Nature 401:877–884.

65. Osburn ED, Weissman JL, Strickland MS, Bahram M, Stone BW, McBride SG. 2025. Relative abundances of bacterial phyla are strong indicators of community-scale microbial growth rates in soil. Environmental Microbiome 20:131.

66. Weissman JL, Hou S, Fuhrman JA. 2021. Estimating maximal microbial growth rates from cultures, metagenomes, and single cells via codon usage patterns. Proceedings of the National Academy of Sciences 118:e2016810118.

67. Signorelli M, Vinciotti V, Wit EC. 2016. NEAT: an efficient network enrichment analysis test. BMC Bioinformatics 17:352.

68. Csardi G, Nepusz T. 2006. The igraph software package for complex network research. InterJournal Complex Systems:1695.

69. Parks DH, Chaumeil P-A, Mussig AJ, Rinke C, Chuvochina M, Hugenholtz P. 2026. GTDB release 10: a complete and systematic taxonomy for 715 230 bacterial and 17 245 archaeal genomes. Nucleic Acids Res 54:D743–D754.

70. Morrissey EM, Mau RL, Schwartz E, Caporaso JG, Dijkstra P, Gestel N van, Koch BJ, Liu CM, Hayer M, McHugh TA, Marks JC, Price LB, Hungate BA. 2016. Phylogenetic organization of bacterial activity. ISME J 10:2336–2340.

71. Stone BWG, Dijkstra P, Finley BK, Fitzpatrick R, Foley MM, Hayer M, Hofmockel KS, Koch BJ, Li J, Liu XJA, Martinez A, Mau RL, Marks J, Monsaint-Queeney V, Morrissey EM, Propster J, Pett-Ridge J, Purcell AM, Schwartz E, Hungate BA. 2023. Life history strategies among soil bacteria—dichotomy for few, continuum for many. ISME J 1–9.

72. Morrissey EM, Mau RL, Hayer M, Liu X-JA, Schwartz E, Dijkstra P, Koch BJ, Allen K, Blazewicz SJ, Hofmockel K, Pett-Ridge J, Hungate BA. 2019. Evolutionary history constrains microbial traits across environmental variation. Nat Ecol Evol 3:1064–1069.

73. Fierer N, Bradford MA, Jackson RB. 2007. Toward an ecological classification of soil bacteria. Ecology 88:1354–1364.

74. Wattenburger CJ, Wang E, Buckley DH. 2025. Dynamics of bacterial growth, and life-history tradeoffs, explain differences in soil carbon cycling due to land-use. ISME Communications 5:ycaf014.

75. Dutilh BE, Varsani A, Tong Y, Simmonds P, Sabanadzovic S, Rubino L, Roux S, Muñoz AR, Lood C, Lefkowitz EJ, Kuhn JH, Krupovic M, Edwards RA, Brister JR, Adriaenssens EM, Sullivan MB. 2021. Perspective on taxonomic classification of uncultivated viruses. Current Opinion in Virology 51:207–215.

76. Howard-Varona C, Hargreaves KR, Abedon ST, Sullivan MB. 2017. Lysogeny in nature: mechanisms, impact and ecology of temperate phages. ISME J 11:1511–1520.

77. Williamson KE, Radosevich M, Smith DW, Wommack KE. 2007. Incidence of lysogeny within temperate and extreme soil environments. Environmental Microbiology 9:2563–2574.

78. Ghosh D, Roy K, Williamson KE, White DC, Wommack KE, Sublette KL, Radosevich M. 2008. Prevalence of Lysogeny among Soil Bacteria and Presence of 16S rRNA and trzN Genes in Viral-Community DNA. Applied and Environmental Microbiology 10.1128/AEM.01435-07.

79. Martin C, Emerson JB, Roux S, Anantharaman K. 2025. A call for caution in the biological interpretation of viral auxiliary metabolic genes. Nat Microbiol 10:2122–2129.

80. Tong D, Ma B, Hu L, Li Y, Dahlgren RA, Xu J. 2025. Soil Virus Life-Strategy Conversion and Implications for Ecosystem and Soil Functions. Global Change Biology 31:e70460.

81. Silpe JE, Bassler BL. 2019. A Host-Produced Quorum-Sensing Autoinducer Controls a Phage Lysis-Lysogeny Decision. Cell 176:268–280.e13.

82. Jansson JK, Wu R. 2023. Soil viral diversity, ecology and climate change. 5. Nat Rev Microbiol 21:296–311.

83. Starr EP, Nuccio EE, Pett-Ridge J, Banfield JF, Firestone MK. 2019. Metatranscriptomic reconstruction reveals RNA viruses with the potential to shape carbon cycling in soil. PNAS 116:25900–25908.

84. Hillary LS, Adriaenssens EM, Jones DL, McDonald JE. 2022. RNA-viromics reveals diverse communities of soil RNA viruses with the potential to affect grassland ecosystems across multiple trophic levels. ISME Commun 2:34.

