## Supplementary Information for "Virome DNA stable isotope probing reveals diverse active soil virus communities across ecosystem contexts"

**Supplementary Methods**

*Soil Physicochemical Analyses*

Water content was determined by mass loss after drying at 105°C and soil pH was determined in a 1:1 soil:DI H_2_O slurry. Soil water holding capacity was measured by wetting soils to field capacity, then measuring water content by mass loss after drying at 105°C for 24 h. We measured extractable N and microbial biomass C using a modified chloroform fumigation-extraction [1] with 0.05M K_2_SO_4_ (1:5 soil:solution). Extracts were measured for C and N on a Shimadzu TOC analyzer (Shimadzu Corp., Kyoto, Japan). Soil total C and N were measured on dried and ground samples on an Elementar soliCube elemental analyzer (Elementar, Langenselbold, Germany). Microbial activity (*i.e*., CO_2_ respiration rate) was measured using a static incubation method [2]. Soil physicochemical data is provided on Table 1.

*DNA extraction and 16S rRNA gene amplicon sequencing*

DNA was extracted from 250 mg of each soil using a Qiagen Powersoil Pro kit (Qiagen, Hilden, Germany). To characterize soil prokaryote communities, we conducted Illumina amplicon sequencing of the V4-V5 region of the 16S rRNA gene using the 515F/926R primer pair [3–5]. PCR reactions contained 10 μl NEB OneTaq Hot Start PCR master mix (New England Biolabs, Ipswich, MA, USA), 2 μl of 1:10 diluted DNA template, 0.2 μM forward and reverse primer, and nuclease-free H_2_O to 25 μl. We also amplified blanks to detect possible contamination. Thermocycling conditions were as follows: 30 s at 94℃ followed by 25 cycles of 45 s at 94℃, 60 s at 50℃, and 90 s at 72℃, with a 10 min final extension at 72℃. Amplicons and negative controls were visualized on a 1% agarose gel to verify successful PCR. Then, amplicons were pooled in equimolar ratios and purified using the Qiagen QIAquick PCR purification kit (Qiagen, Valencia, CA, USA). The purified sequence libraries were then sequenced on an Illumina MiSeq with 250 bp paired-end reads. The raw amplicon sequence reads are available at NCBI accession# PRJNA1471083.

Raw sequences were bioinformatically processed using the DADA2 pipeline [6] and taxonomy assigned to the resulting amplicon sequence variants (ASVs) using the IDTAXA algorithm [7] trained on the SILVA SSU database (version 138.2) [8]. Prior to downstream analyses, we filtered eukaryote contaminants from the 16S dataset, as well as sequences classified as chloroplasts, mitochondria, and sequences unclassified to a prokaryote phylum. Prior to analyses, we rarefied the prokaryote dataset to 12,135 sequences per sample.

**Supplementary Tables and Figures**

**Supplementary Table S1**

| **Sequencing Project ID** | **Library Name** | **BioProject** | **BioSample** | **Project Accession** | **Run Accession** |
| --- | --- | --- | --- | --- | --- |
| 1583100 | PHBTW | PRJNA1450189 | SAMN57148267 | SRP690350 | SRR38009072 |
| 1583101 | PHBTX | PRJNA1450139 | SAMN57147623 | SRP690348 | SRR38009070 |
| 1583102 | PHBTY | PRJNA1450184 | SAMN57147442 | SRP690351 | SRR38009073 |
| 1583103 | PHBTZ | PRJNA1450165 | SAMN57147254 | SRP690352 | SRR38009078 |
| 1583104 | PHBUA | PRJNA1450162 | SAMN57147258 | SRP690353 | SRR38009082 |
| 1583105 | PHBUB | PRJNA1450134 | SAMN57147624 | SRP690355 | SRR38009088 |
| 1583106 | PHBUC | PRJNA1450164 | SAMN57145987 | SRP690354 | SRR38009087 |
| 1583107 | PHBUG | PRJNA1450156 | SAMN57145983 | SRP690356 | SRR38009093 |
| 1583108 | PHBUH | PRJNA1450152 | SAMN57147260 | SRP690288 | SRR38007226 |
| 1583110 | PHBUO | PRJNA1450133 | SAMN57147517 | SRP690290 | SRR38007236 |
| 1583111 | PHBUP | PRJNA1450177 | SAMN57148238 | SRP690291 | SRR38007237 |
| 1583112 | PHBUS | PRJNA1450150 | SAMN57147255 | SRP690292 | SRR38007238 |
| 1583113 | PHBUT | PRJNA1450128 | SAMN57147626 | SRP690293 | SRR38007239 |
| 1583114 | PHBUU | PRJNA1450155 | SAMN57147485 | SRP690299 | SRR38007259 |
| 1583115 | PHBUW | PRJNA1450153 | SAMN57147256 | SRP690294 | SRR38007240 |
| 1583117 | PHBUY | PRJNA1450188 | SAMN57147515 | SRP690295 | SRR38007241 |
| 1583118 | PHBUZ | PRJNA1450129 | SAMN57147516 | SRP690298 | SRR38007258 |
| 1583120 | PHBWB | PRJNA1450149 | SAMN57145981 | SRP690296 | SRR38007245 |
| 1583121 | PHBWC | PRJNA1450182 | SAMN57147625 | SRP690300 | SRR38007260 |
| 1583122 | PHBWG | PRJNA1450171 | SAMN57147441 | SRP690305 | SRR38007274 |
| 1583123 | PHBWH | PRJNA1450163 | SAMN57147257 | SRP690302 | SRR38007267 |
| 1583125 | PHBWO | PRJNA1450187 | SAMN57148230 | SRP690303 | SRR38007268 |
| 1583126 | PHBWP | PRJNA1450185 | SAMN57147439 | SRP690304 | SRR38007269 |
| 1583127 | PHBWS | PRJNA1450178 | SAMN57147259 | SRP690308 | SRR38007277 |
| 1583128 | PHBWT | PRJNA1450154 | SAMN57145992 | SRP690306 | SRR38007275 |
| 1583129 | PHBWU | PRJNA1450127 | SAMN57147438 | SRP690307 | SRR38007276 |
| 1583130 | PHBWW | PRJNA1450175 | SAMN57147514 | SRP690309 | SRR38007278 |
| 1583131 | PHBWX | PRJNA1450157 | SAMN57147252 | SRP690310 | SRR38007279 |
| 1583132 | PHBZB | PRJNA1450168 | SAMN57148228 | SRP690313 | SRR38008207 |
| 1583134 | PHBZG | PRJNA1450180 | SAMN57148265 | SRP690314 | SRR38008210 |
| 1583135 | PHBZH | PRJNA1450170 | SAMN57145979 | SRP690906 | SRR38047030 |
| 1583136 | PHBZN | PRJNA1450142 | SAMN57145993 | SRP690317 | SRR38008213 |
| 1583137 | PHBZO | PRJNA1450136 | SAMN57145994 | SRP690315 | SRR38008212 |
| 1583138 | PHBZP | PRJNA1450172 | SAMN57145986 | SRP690316 | SRR38008214 |
| 1583139 | PHBZS | PRJNA1450145 | SAMN57147253 | SRP690319 | SRR38008242 |
| 1583140 | PHBZT | PRJNA1450148 | SAMN57145980 | SRP690320 | SRR38008243 |
| 1583141 | PHBZU | PRJNA1450125 | SAMN57145982 | SRP690321 | SRR38008244 |
| 1583142 | PHBZW | PRJNA1450126 | SAMN57147486 | SRP690323 | SRR38008246 |
| 1583143 | PHBZX | PRJNA1450174 | SAMN57147488 | SRP690322 | SRR38008245 |
| 1583144 | PHBZY | PRJNA1450141 | SAMN57145985 | SRP690325 | SRR38008254 |
| 1583145 | PHBZZ | PRJNA1450140 | SAMN57147513 | SRP690327 | SRR38008270 |
| 1583146 | PHCAA | PRJNA1450169 | SAMN57145978 | SRP690358 | SRR38009131 |
| 1583147 | PHCAB | PRJNA1450146 | SAMN57145984 | SRP690328 | SRR38008271 |
| 1583148 | PHCAC | PRJNA1450161 | SAMN57147627 | SRP690329 | SRR38008272 |
| 1583150 | PHCAH | PRJNA1450183 | SAMN57147487 | SRP690330 | SRR38008273 |
| 1583151 | PHCAN | PRJNA1450167 | SAMN57147436 | SRP690332 | SRR38008275 |
| 1583152 | PHCAO | PRJNA1450138 | SAMN57145975 | SRP690331 | SRR38008274 |
| 1583153 | PHCAP | PRJNA1450173 | SAMN57148266 | SRP690333 | SRR38008276 |
| 1583155 | PHCAT | PRJNA1450130 | SAMN57145977 | SRP690337 | SRR38008735 |
| 1583156 | PHCAU | PRJNA1450143 | SAMN57148229 | SRP690339 | SRR38009035 |
| 1583158 | PHCAX | PRJNA1450179 | SAMN57148237 | SRP690341 | SRR38009036 |
| 1583159 | PHCAY | PRJNA1450166 | SAMN57147437 | SRP690340 | SRR38009037 |
| 1583160 | PHCAZ | PRJNA1450137 | SAMN57145974 | SRP690344 | SRR38009041 |
| 1583161 | PHCBA | PRJNA1450135 | SAMN57147512 | SRP690346 | SRR38009061 |
| 1583162 | PHCBB | PRJNA1450144 | SAMN57147440 | SRP690343 | SRR38009040 |
| 1583163 | PHCBC | PRJNA1450131 | SAMN57145976 | SRP690349 | SRR38009071 |

Supplementary Table S1: NCBI accession numbers for the raw NovaSeq virus metagenome sequence reads from each sample.

**Supplementary Table S2**

| **sample** | **n_reads** | **n_bp** | **n_contigs** | **contig_bp** | **ctg_N90** | **ctg_L90** | **ctg_N50** | **ctg_L50** | **Soil** |
| --- | --- | --- | --- | --- | --- | --- | --- | --- | --- |
| s17 | 67780522 | 10096461754 | 212147 | 169600905 | 26994 | 1216 | 152293 | 317 | Ag |
| s18 | 69317520 | 10326850925 | 187589 | 158395675 | 22808 | 1364 | 131682 | 323 | Ag |
| s19 | 67997438 | 10129221534 | 289757 | 220677895 | 37468 | 1087 | 213276 | 313 | Ag |
| s20 | 67052854 | 9983591711 | 262853 | 210456611 | 32615 | 1205 | 189679 | 319 | Ag |
| s21 | 62076198 | 9237874714 | 203782 | 161744810 | 26562 | 1213 | 146679 | 316 | Ag |
| s22 | 82766670 | 12321090059 | 173531 | 133920558 | 23604 | 1129 | 126006 | 315 | Ag |
| s23 | 59207718 | 8810077528 | 220365 | 185350903 | 27130 | 1331 | 156447 | 324 | Ag |
| s24 | 69939146 | 10412190082 | 204901 | 164502558 | 26417 | 1219 | 147533 | 319 | Ag |
| s25 | 89346834 | 13311951248 | 832150 | 672940878 | 99366 | 1131 | 605094 | 324 | Forest |
| s26 | 120873438 | 18015949799 | 693339 | 523925500 | 87760 | 998 | 513641 | 312 | Forest |
| s27 | 100356630 | 14947755544 | 772102 | 739133876 | 68483 | 1635 | 528666 | 347 | Forest |
| s28 | 90637818 | 13500437541 | 685310 | 655688104 | 61098 | 1631 | 470390 | 346 | Forest |
| s29 | 117833692 | 17550268811 | 589082 | 476292399 | 65882 | 1171 | 425293 | 318 | Forest |
| s30 | 99028874 | 14733090023 | 767420 | 782877557 | 83564 | 1983 | 495923 | 359 | Forest |
| s31 | 114885336 | 17115680386 | 1182186 | 919183672 | 148891 | 1047 | 869812 | 319 | Forest |
| s32 | 104327174 | 15544707126 | 830795 | 647569388 | 103180 | 1051 | 612952 | 319 | Forest |
| s33 | 67866182 | 10109176988 | 248585 | 185507910 | 30233 | 1204 | 171679 | 305 | Ag |
| s34 | 73200710 | 10905183298 | 238518 | 184832539 | 29045 | 1229 | 166762 | 311 | Ag |
| s35 | 67766352 | 10094564174 | 286960 | 212284559 | 39200 | 1021 | 213544 | 311 | Ag |
| s36 | 56058998 | 8347417759 | 266772 | 201646191 | 36452 | 1066 | 196971 | 314 | Ag |
| s37 | 66071124 | 9832107449 | 227154 | 186060509 | 29773 | 1275 | 161388 | 321 | Ag |
| s39 | 66734440 | 9931506652 | 271308 | 212703532 | 36340 | 1164 | 197396 | 316 | Ag |
| s40 | 56571616 | 8424492892 | 230167 | 184575124 | 29513 | 1219 | 166371 | 318 | Ag |
| s41 | 126456388 | 18842690104 | 1196663 | 978339166 | 138803 | 1171 | 865055 | 324 | Forest |
| s42 | 112823992 | 16814825020 | 1027369 | 865824885 | 117456 | 1236 | 736355 | 329 | Forest |
| s43 | 103935090 | 15487932752 | 751198 | 715331391 | 65907 | 1649 | 515276 | 343 | Forest |
| s44 | 98301048 | 14647162557 | 688248 | 684381206 | 58294 | 1793 | 464735 | 351 | Forest |
| s46 | 95374330 | 14208665925 | 794343 | 697439000 | 82689 | 1365 | 562453 | 332 | Forest |
| s47 | 82268224 | 12257951220 | 916735 | 711272589 | 125629 | 1026 | 676820 | 324 | Forest |
| s48 | 98968160 | 14748806423 | 1063491 | 825585040 | 141769 | 1041 | 784475 | 321 | Forest |
| s49 | 53199936 | 7927746338 | 183273 | 152126875 | 19386 | 1389 | 126252 | 319 | Ag |
| s50 | 71565984 | 10662812829 | 290703 | 218658008 | 39522 | 1064 | 214372 | 313 | Ag |
| s51 | 60912816 | 9076735098 | 193564 | 155904223 | 23474 | 1204 | 140048 | 320 | Ag |
| s52 | 52644976 | 7841804284 | 296319 | 206342992 | 45007 | 893 | 224685 | 308 | Ag |
| s53 | 61919502 | 9222031704 | 211003 | 163272473 | 27896 | 1127 | 154152 | 315 | Ag |
| s54 | 66836238 | 9953417889 | 228075 | 162562701 | 33398 | 959 | 171311 | 308 | Ag |
| s55 | 62795634 | 9348082759 | 241301 | 196388285 | 29472 | 1228 | 173381 | 321 | Ag |
| s56 | 44086528 | 6565477566 | 280614 | 220223874 | 37431 | 1144 | 204464 | 319 | Ag |
| s57 | 91852820 | 13690002524 | 897991 | 730577231 | 108921 | 1129 | 652109 | 327 | Forest |
| s58 | 122058296 | 18189984370 | 1060606 | 920786316 | 109933 | 1336 | 748280 | 331 | Forest |
| s59 | 98619136 | 14696348408 | 632308 | 615167894 | 50352 | 1738 | 430132 | 346 | Forest |
| s60 | 101871966 | 15178764615 | 651978 | 651447968 | 50705 | 1844 | 438805 | 350 | Forest |
| s61 | 104554412 | 15580796152 | 888557 | 772720659 | 90717 | 1324 | 629467 | 332 | Forest |
| s62 | 103249326 | 15383367619 | 934654 | 846935100 | 92891 | 1447 | 654899 | 338 | Forest |
| s63 | 101106964 | 15067482914 | 930268 | 795845604 | 98137 | 1276 | 665187 | 330 | Forest |
| s64 | 106901220 | 15930204851 | 1024987 | 839019908 | 115188 | 1172 | 740959 | 324 | Forest |

Supplementary Table S2: Statistics on the quality-filtered sequence reads and assemblies for each of the virus metagenomes.

Supplementary Table S3

| **Taxonomic Level** | **Ag. Soils** | | **Forest Soils** | |
| --- | --- | --- | --- | --- |
|  | Adj. R^2^ | *P* value | Adj. R^2^ | *P* value |
| Phylum | 0.135 | < 0.001 | 0.101 | < 0.001 |
| Class | 0.135 | < 0.001 | 0.187 | < 0.001 |
| Order | 0.258 | < 0.001 | 0.303 | < 0.001 |
| Family | 0.403 | < 0.001 | 0.328 | < 0.001 |

Supplementary Table S3: Proportions of variation in vOTU ^13^C EAF explained by host taxonomic levels. Adjusted R^2^ values and p values are from linear models. At each taxonomic level, taxa with fewer than 5 representatives in the dataset were removed prior to analysis.

**Supplementary Figure S1**

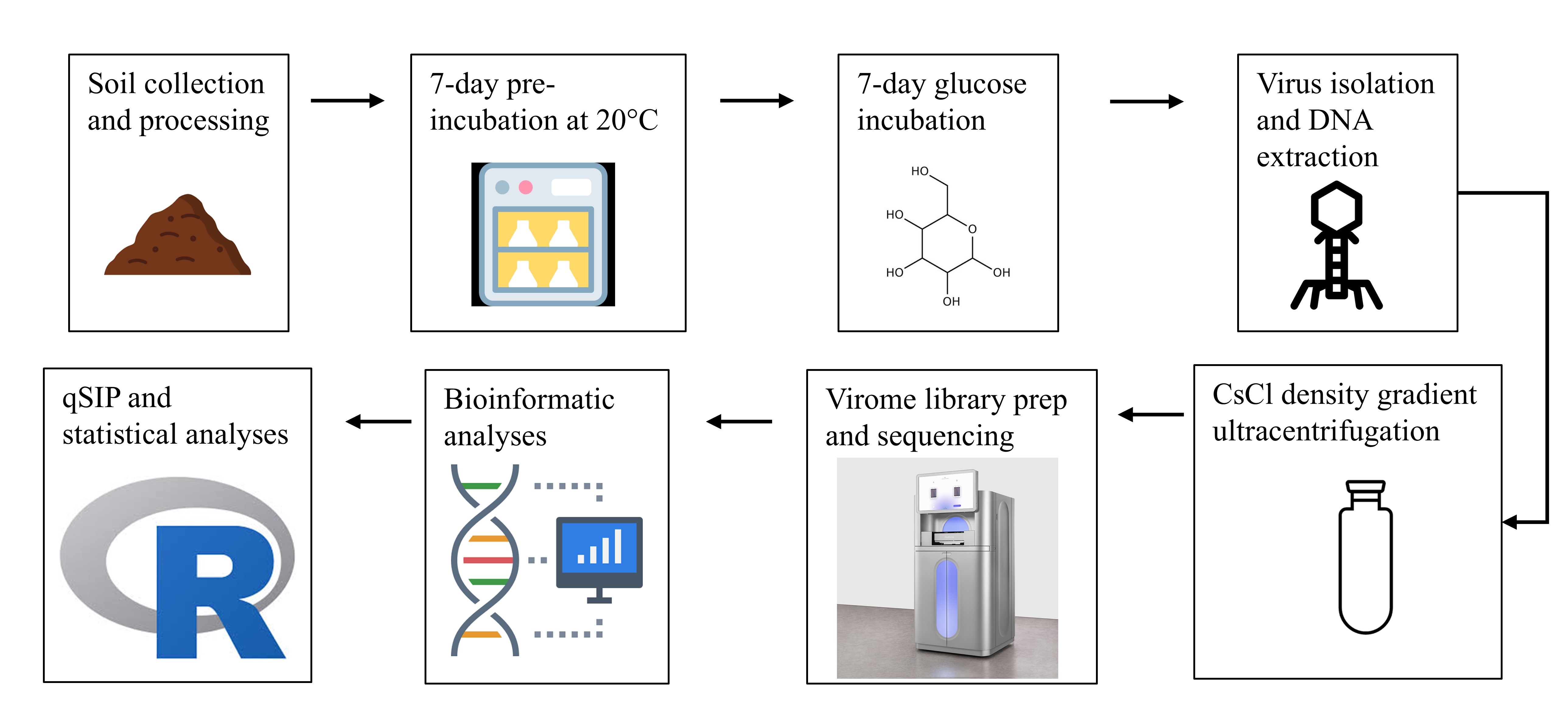

Supplementary Fig. S1: Schematic overview of the virome DNA-SIP experimental procedures.

**Supplementary Figure S2**

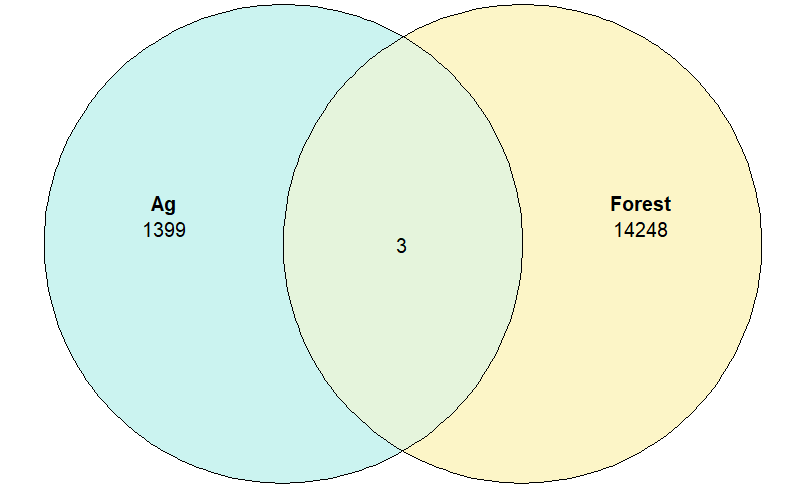

Supplementary Fig. S2: Unique and shared vOTUs between the agricultural and forest soils.

**Supplementary Figure S3**

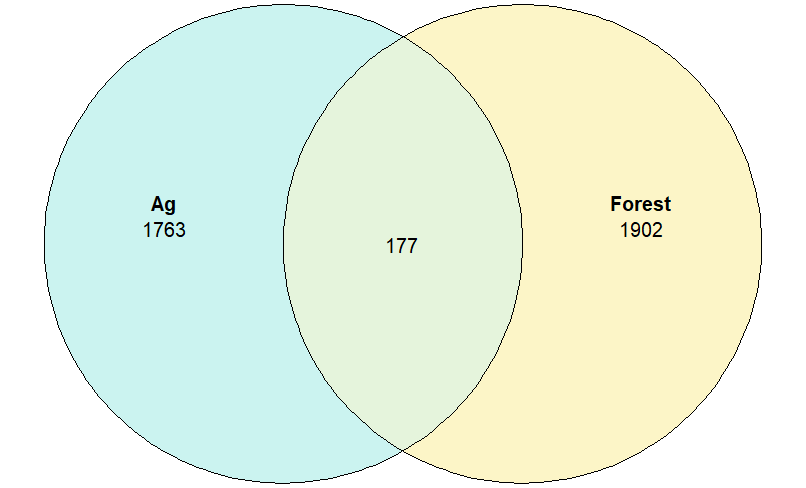

Supplementary Fig. S3: Unique and shared prokaryote ASVs between the agricultural and forest soils.

**Supplementary Figure S4**

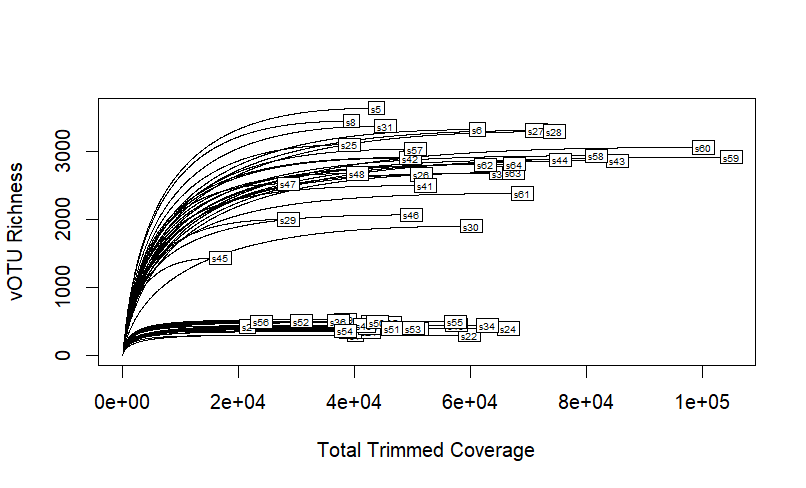

Supplementary Fig. S4: Alpha rarefaction curves showing vOTU richness in each sample as a function of sequencing effort (total coverage) in each sample.

**Supplementary Fig. S5**

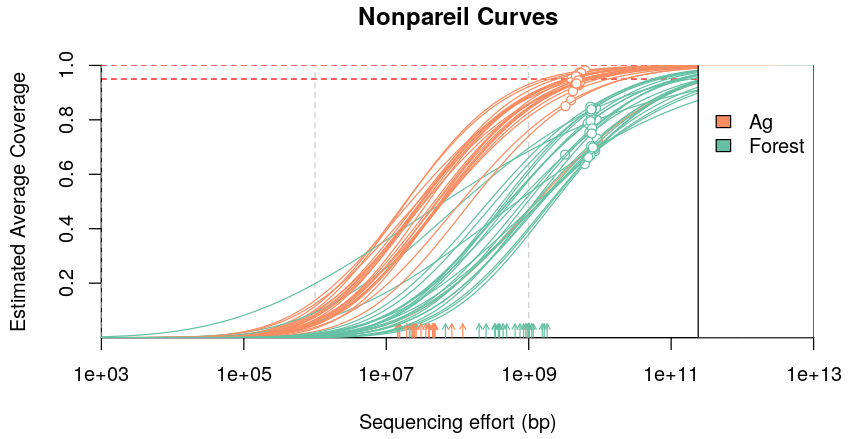

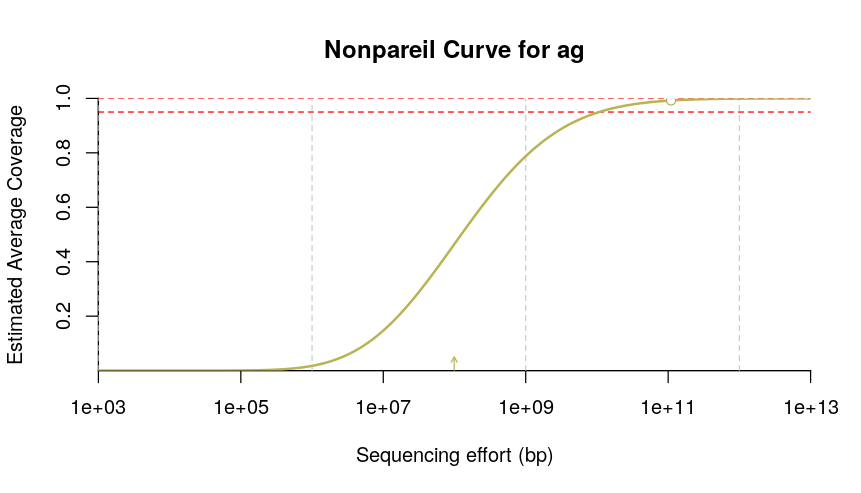

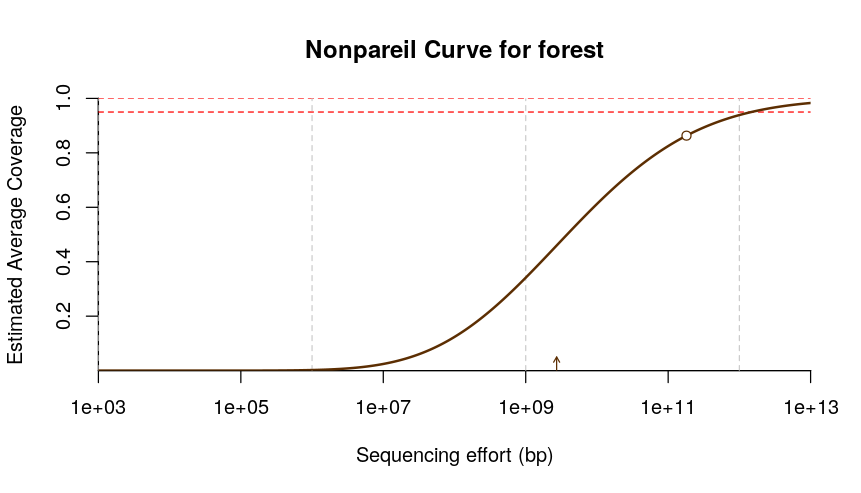

Supplementary Fig. S5: Nonpareil coverage curves showing estimated average coverage as a function of sequence depth. Circles on the curves indicate the actual position of each sample on its coverage curve. Coverage was generally higher in the agricultural soils (mostly >90% individually and essentially 100% for the co-assembly) than in the forest soils (mostly 60-80% individually and ~85% for the co-assembly).

**Supplementary Figure S6**

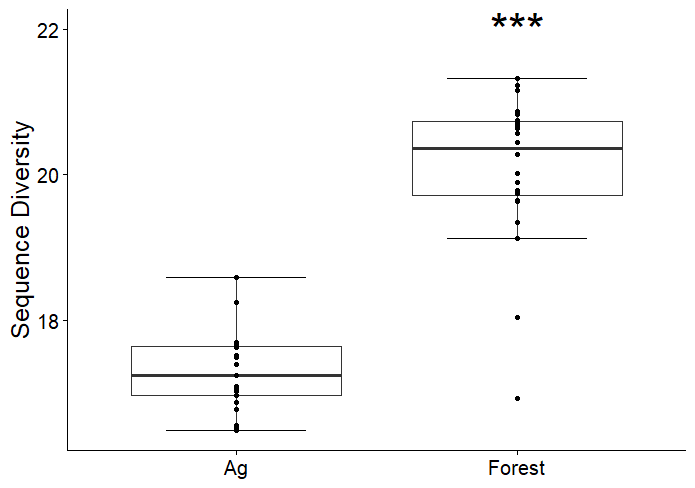

Supplementary Fig. S6: Nonpareil analysis of sequence-level diversity. Asterisks indicate significantly higher diversity (one-way ANOVA) at the following significance level: *** *p*<0.001

**Supplementary Figure S7**

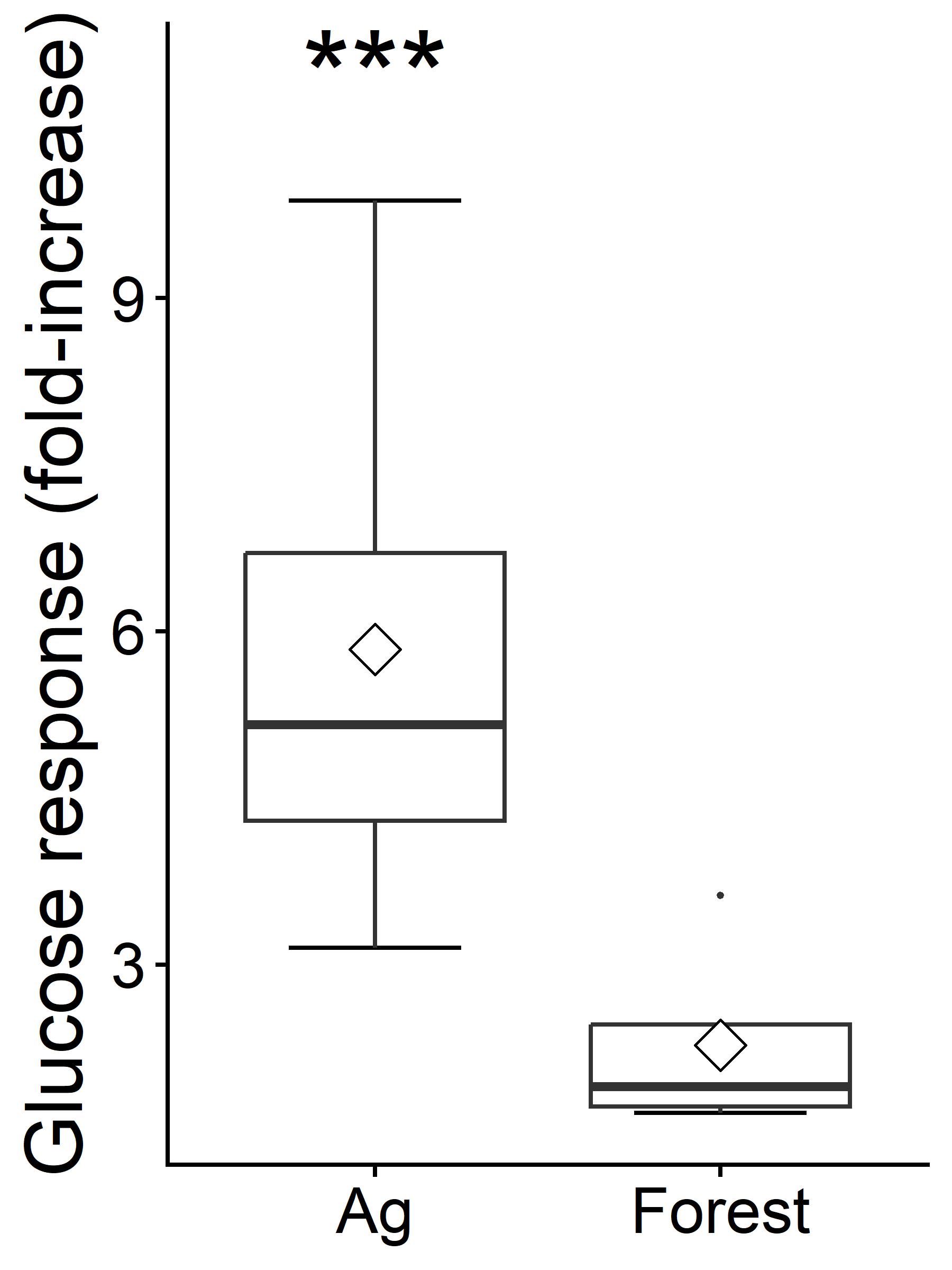

Supplementary Fig. S7: Respiratory responses to the glucose addition. The agricultural soil had ~2.6-fold greater respiratory responses to the glucose addition (relative to no-glucose control respiration) on average.

**Supplementary Figure S8**

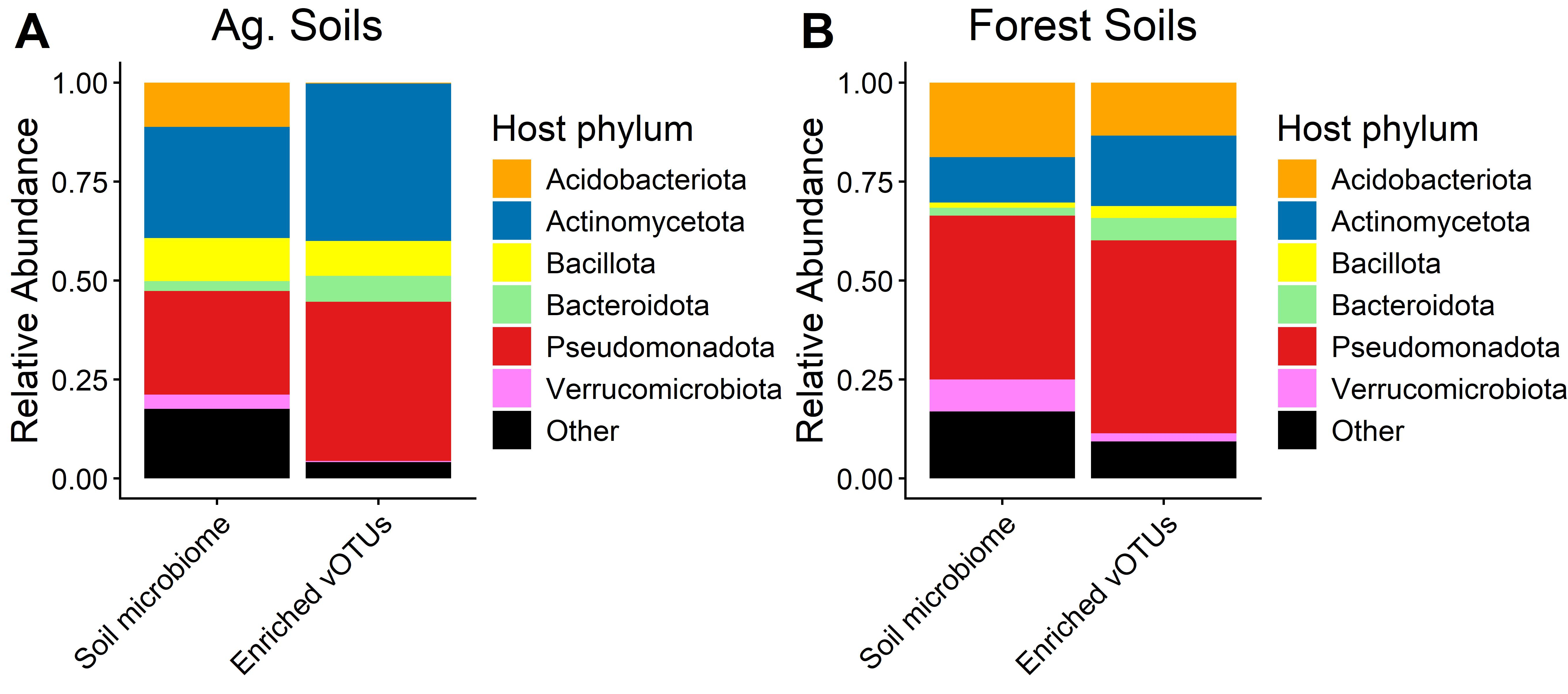

Supplementary Fig. S8: Taxonomic composition of the underlying soil microbiomes and the 13C-enriched (active) vOTU microbial hosts in the agricultural (A) and forest (B) soils.

**Supplementary Figure S9**

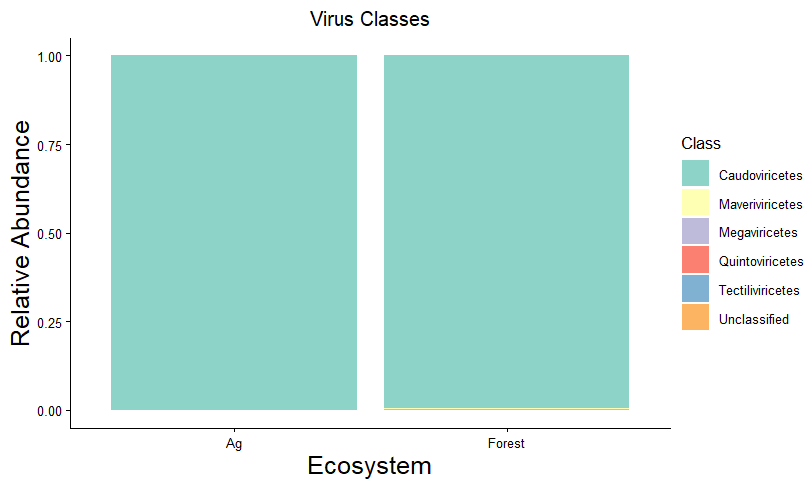

Supplementary Fig. S9: Relative abundances of virus classes in the agricultural and forest soils. Average virus class relative abundances across the dataset were as follows: Caudoviricetes (99.7%), Maveriviricetes (0.18%), Megaviricetes (0.0065%), Quintoviricetes (0.0011%), Tectiliviricetes (0.036%), Unclassified (0.08%).

**Supplementary Figure S10**

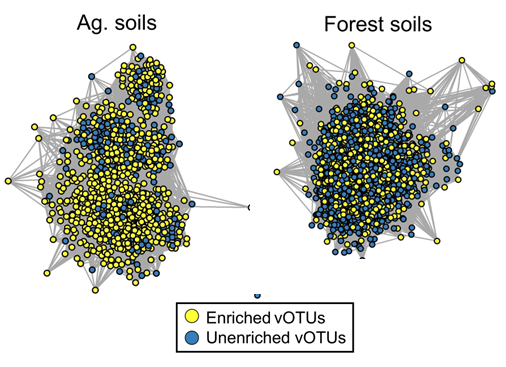

Supplementary Fig. S10: Comparisons of the ^13^C enriched and unenriched vOTU genomes from the two soil types. Shown are gene-sharing networks for both soil types, with ^13^C-enriched and unenriched genomes (the network nodes) indicated. Hypergeometric tests indicated that enriched vOTU genomes from both soil types occupied distinct clusters in the gene-sharing networks, i.e., enriched and unenriched genomes had fewer links than expected by chance (p < 0.001).

**References**

1.	Witt C et al. A rapid chloroform-fumigation extraction method for measuring soil microbial biomass carbon and nitrogen in flooded rice soils. *Biology and Fertility of Soils* 2000;**30**:510–519. https://doi.org/10.1007/s003740050030

2.	Bradford MA, Fierer N, Reynolds JF. Soil carbon stocks in experimental mesocosms are dependent on the rate of labile carbon, nitrogen and phosphorus inputs to soils. *Funct Ecol* 2008;**22**:964–974. https://doi.org/10.1111/j.1365-2435.2008.01404.x

3.	Parada AE, Needham DM, Fuhrman JA. Every base matters: assessing small subunit rRNA primers for marine microbiomes with mock communities, time series and global field samples. *Environ Microbiol* 2016;**18**:1403–1414. https://doi.org/10.1111/1462-2920.13023

4.	Quince C et al. Removing Noise From Pyrosequenced Amplicons. *BMC Bioinformatics* 2011;**12**:38. https://doi.org/10.1186/1471-2105-12-38

5.	Walters W et al. Improved Bacterial 16S rRNA Gene (V4 and V4-5) and Fungal Internal Transcribed Spacer Marker Gene Primers for Microbial Community Surveys. *mSystems* 2016;**1**:e00009-15. https://doi.org/10.1128/mSystems.00009-15

6.	Callahan BJ et al. DADA2: High-resolution sample inference from Illumina amplicon data. *Nat Methods* 2016;**13**:581–583. https://doi.org/10.1038/nmeth.3869

7.	Murali A, Bhargava A, Wright ES. IDTAXA: a novel approach for accurate taxonomic classification of microbiome sequences. *Microbiome* 2018;**6**:140. https://doi.org/10.1186/s40168-018-0521-5

8.	Quast C et al. The SILVA ribosomal RNA gene database project: improved data processing and web-based tools. *Nucleic Acids Res* 2013;**41**:D590–D596. https://doi.org/10.1093/nar/gks1219
